# Microenvironment Drives Reentrant Condensation of Aβ40

**DOI:** 10.1101/2024.04.27.591429

**Authors:** Susmita Sarkar, Jagannath Mondal

**Affiliations:** Tata Institute of Fundamental Research Hyderabad 36/P Gopanapally village, Hyderabad, Telangana, India 500046

## Abstract

Within the framework of liquid-liquid phase separation (LLPS), biomolecular condensation orchestrates vital cellular processes and its dysregulation is implicated in severe pathological conditions. Recent studies highlight the role of intrinsically disordered proteins (IDPs) in LLPS, yet the influence of microenvironmental factors has remained a puzzling factor. Here, via computationally simulating the impact of solution conditions on LLPS behavior of neurologically pathogenic IDP Aβ40, we chanced upon a salt-driven reentrant condensation phenomenon, wherein Aβ40 aggregation increases with low salt concentrations (25-50 mM), followed by a decline with further salt increments.. An exploration into the thermodynamic and kinetic signatures of reentrant condensation unveils a nuanced interplay between protein electrostatics and ionic strength as potential drivers. Notably, the charged residues of the N-terminus exhibit a non-monotonic response to salt screening, intricately linked to the recurrence of reentrant behavior in hydrophobic core-induced condensation. Intriguingly, our findings also unveil the reappearance of similar reentrant condensation phenomena under varying temperature conditions. Collectively, our study illuminates the profoundly context-dependent nature of Aβ40’s liquid-liquid phase separation behavior, extending beyond its intrinsic molecular framework, where microenvironmental cues wield significant influence over its aberrant functionality.

## Introduction

LLPS has surfaced as a pivotal organizational principle in the realm of biology where cellular components such as proteins and nucleic acids undergo condensation into liquid droplets, facilitating the formation of membraneless subcellular compartments that orchestrate essential biological processes^1–3^. Beyond cellular compartmentalization, biomolecular condensation serves as a crucial mechanism guiding various biological phenomena, encompassing sensory perception, RNA metabolism, chromatin dynamics, stress response, and noise modulation. Dysregulation of biomolecular condensates has been implicated in the onset of severe pathological conditions, including neurodegenerative disorders and cancer^3,4^. This thermodynamic phenomenon entails the reduction of free energy as proteins undergo demixing, partitioning into a protein-rich condensed phase and a protein-deficient dilute phase^5^. The attainment of equilibrium between biomolecular partitioning into these phases poses a fundamental inquiry into the driving forces governing intermolecular interactions and the factors modulating LLPS dynamics.

A surge of recent investigations has shown that IDPs are well suited to be involved in phase separation^6–13^. The specific molecular grammar involving the interrelationship of the primary sequence of a biopolymer and its ability to undergo phase separation is quite well explored in many previous studies^14–16^. However, some recent studies also hinted that even beyond the unique nature of protein sequence, LLPS is crucially guided by the specific solution conditions^13,17–26^. Different external stimuli like variation in biomolecular concentration, alteration of salt concentration, temperature, pressure, pH, and presence of some biomolecules involving chaperones or ATP can significantly modulate protein LLPS. Despite active research in this field at present, a comprehensive understanding on the role of such microenvironmental factors as molecular driving forces for LLPS is currently elusive. Interactions among biological macromolecules in water are typically intricate and largely influenced by various environmental factors which eventually contribute to diverse phase behavior and play a pivotal role in the onset of several protein aggregation-related diseases.

In this current article, we report an interesting mode of microenvironment-induced alteration of condensation propensity of Aβ40, an IDP associated with Alzheimer’s Disease. First we present computer simulation that definitively showcases LLPS-like characteristics within the aggregates of Aβ40. Next, as an interesting outcome, by computationally simulating the phase separation behaviors of Aβ40 via variation of a wide range of ionic strength in solution condition, our investigation identifies a phenomenon of reentrant condensation of Aβ40 at low salt concentration: the extent of aggregation of Aβ40 is found to rise with increase of salt concentration from 25 mM to 50 mM, while further increase of salt concentration results in gradual reduction in aggregation propensity. This observation is found to be statistically robust and uncovers the complex interplay of thermodynamic forces driven by ionic-strength altered protein electrostatics which eventually modulate the key hydrophobic interactions responsible for amyloid aggregation. Finally, we show that the observation of non-monotonic aggregation is not restricted to only variation of salt. In fact, as would be expanded towards the end of this article, we came across the observed reentrant condensation of Aβ40 across variation of solution temperature as well.

## Results and discussion

### LLPS-like phase separation of Aβ40 captured in computer simulation

Our investigation commenced with the computational modeling of the early-stage aggregation of Aβ40, as depicted schematically in Figure 1A, utilizing the Martini-3 coarse-grained model^27^. Martini, a semi-quantitative model renowned for its efficiency in simulating protein aggregation, strikes a balance between resolution and computational affordability when compared to the computationally intensive all-atom force fields. Notably, Martini distinguishes itself from other recently developed coarse-grained models^28–30^ by its explicit representation of water and salts, albeit at a reduced resolution relative to all-atom models, a feature pivotal for our current investigation (Figure 1A). Protein, water and ions are all described using Martini-3 coarse-grained model and was properly benchmarked against atomistic simulations (see Method and Figure S1-2).

**Figure 1:**
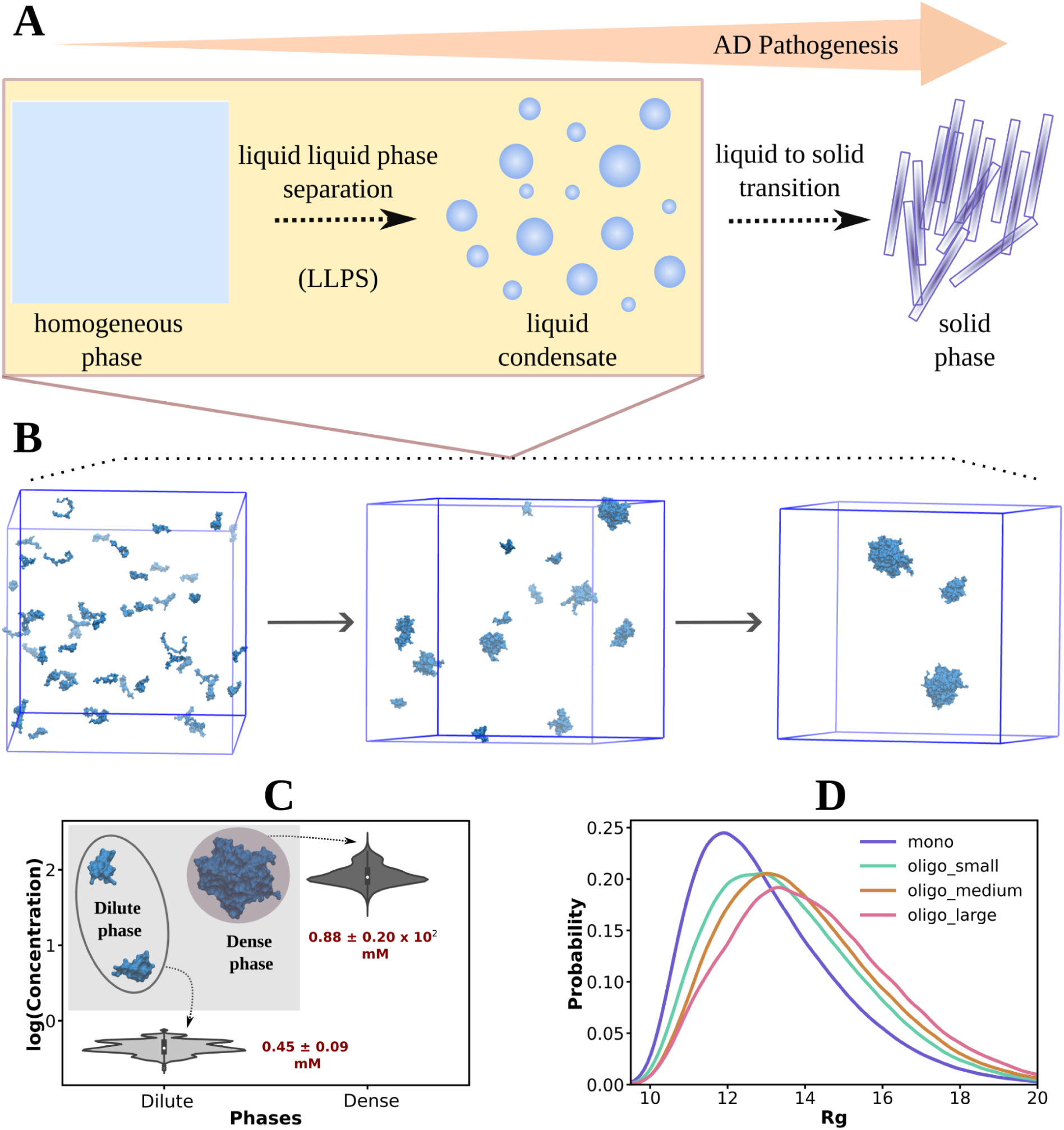
LLPS-like properties of Aβ40 protein in aqueous saline solution. A. Schematic representation of the major steps of AD pathogenesis involving LLPS forming liquid condensates followed by liquid to solid transition. B. Representative simulation snapshots of Aβ40 protein (750 μM in 50 mM NaCl salt at 300 K temperature) starting from initial state involving randomly dispersed protein chains followed by intermediate stage showing some extent of protein oligomerization and finally near final stage involving the formation of larger oligomers. Figure C. represents the probability distribution of protein concentrations (in the form of logarithmic function) in dilute and dense phases in violin plots. The mean and standard deviations are highlighted in red text. Inset shows the schematic representation of the dilute and dense phases involving monomers to smaller oligomers (involving number of protein chains < 6) and larger oligomers (≥ 6 number of protein chains) respectively. Figure D. represents probability distribution of Rg of protein chains in monomeric state and differently sized olimers (small (number of protein chains ≤ 3), intermediate (number of protein chains ≤ 6) and large (number of protein chains > 6)).

We investigated the collective interactions among a set of randomly dispersed chains of Aβ40 protein in an aqueous solution containing 50 mM NaCl at an ambient temperature of 300 K. The system involving randomly dispersed protein chains was further solvated by addition of water and salt in a cubic box of dimensions 48×48×48 nm^3^. This configuration corresponded to a protein concentration of approximately 750 μM, a value that lies within the range of previous experimental studies ^31–33^. Over the course of multiple replicas of 2.5 μs each, the molecular dynamics simulations captured the spontaneous formation of smaller aggregates, representing an intermediate state emerging from the initial state of randomly dispersed protein monomers (Figure 1B). Subsequently, these aggregates underwent further evolution, progressively increasing in size within the system (Figure 1B), indicative of the formation of condensates from a homogeneous phase.

To assess whether the aggregates formed during the simulations possess LLPS-like properties, concentrations of protein in the dilute (involving number of protein chains < 6^24^) and dense phases (≥ 6^24^ number of protein chains) following the equation represented as follows (equation 1)

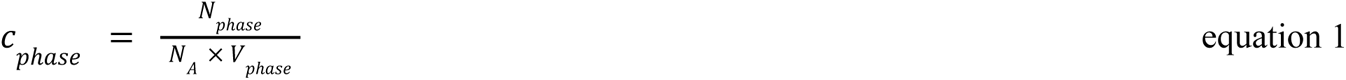

Where c_phase_ is the protein concentration in a phase (either dilute or dense), N_phase_ denotes the number of protein chains occurring in that phase. N_A_ is the Avogadro’s number and the volume occupied by the phase is designated as V_phase_. Volume of the dense phase (V_dense_) is determined using the following equation^34^ (equation 2)

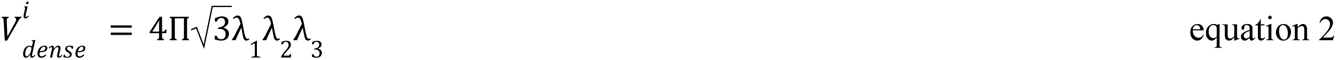

Where V^i^_dense_ denotes the i^th^ droplet volume and the eigenvalues of the gyration tensor of the aggregates has been designated by λ_1_, λ_2_ and λ_3_. The volume of the dilute phase is estimated by considering the volume of the box other than that belonging to the dense phase (equation 3).

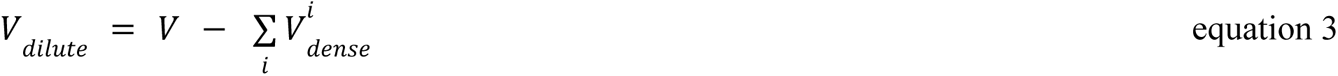

Where V_dilute_ is the volume of the dilute phase and the total volume of the box is represented by V.

Figure S3 represents that each of the replicates of the simulation reaches well in equilibrium during the course of time (1.5 to 2.5 μs). The analysis of protein concentration was performed over the combined trajectories of the last 1 μs from each of the three replica simulations. Figure 1C indicates a nearly three orders of magnitude difference in protein concentrations between the dilute and dense phases. This stark contrast in concentrations is a characteristic hallmark of liquid-liquid phase separation (LLPS), affirming that the current setup effectively captures the observed features of LLPS.

To characterize the structural attributes of the protein chains within the condensates, we compared the average radius of gyration (R_g_) of protein chains in their monomeric state with those in differently sized oligomers (small: number of protein chains ≤ 3, intermediate: number of protein chains ≤ 6, and large: number of protein chains > 6). As depicted in Figure 1D, the Aβ40 chains exhibit greater extension in an oligomeric state compared to their randomly dispersed counterparts, with the extent of chain extension increasing proportionally with the size of the oligomers. This observation of chain extension in aggregate aligns well with recent studies conducted on various IDPs and other bio-macromolecules^24,34–36^.

To evaluate the the trend of observed LLPS-like properties at the present protein concentration (750 μM), additional simulations were conducted at both lower (500 μM) and higher (1000 μM) protein concentrations (see Table 1 for simulation system details) while maintaining a constant salt concentration (50 mM) and temperature (300 K). In all cases, coexistence of dilute and dense phases was observed (see Figure S4), with the extent of phase separation gradually increasing as the protein concentration was raised from 500 to 750 to 1000 μM (see Figure S5).

**Table1:**
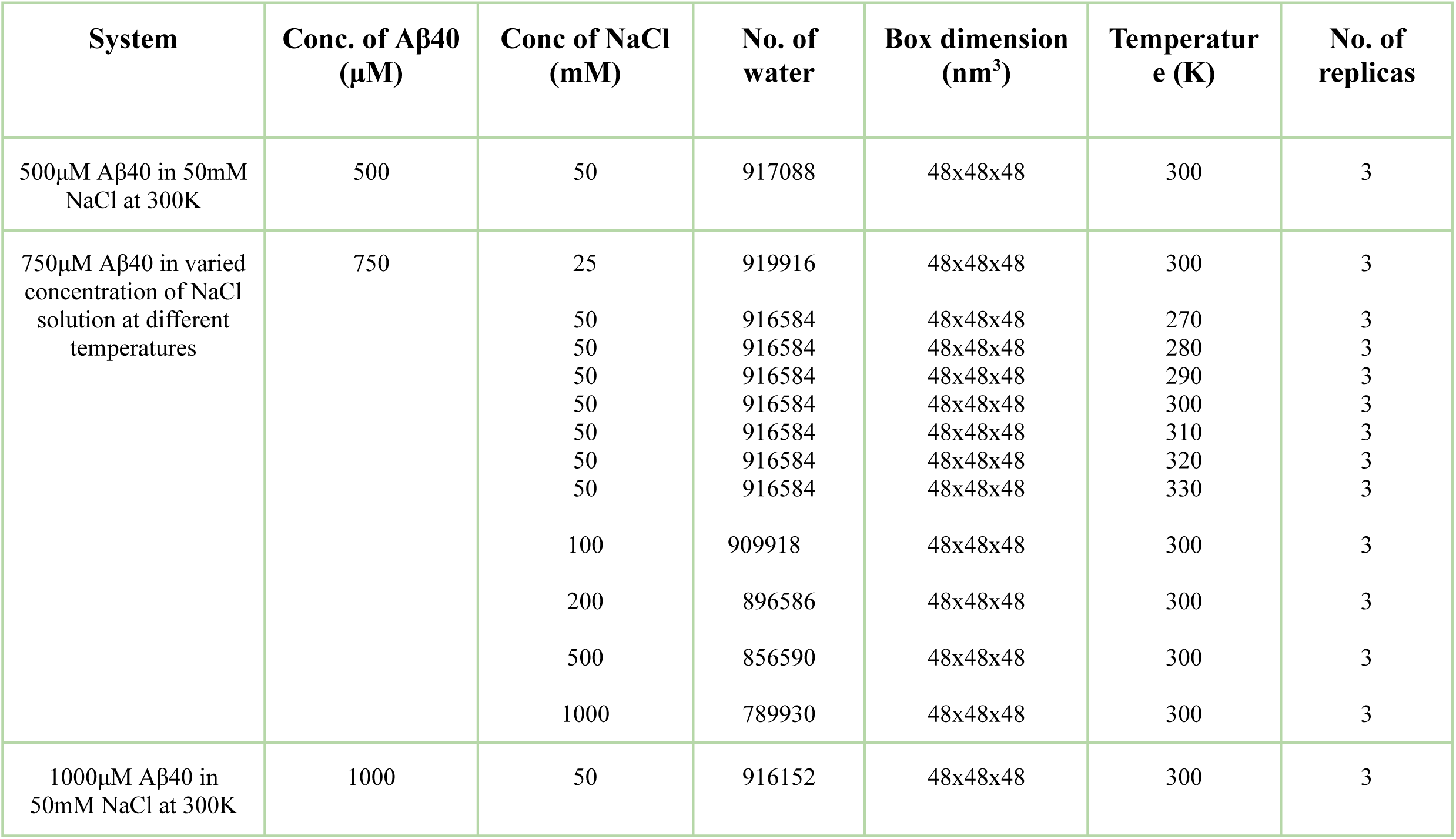
System details of all aggregation simulations performed in this investigation. All constituents were parameterized with Martini-3 coarse-grained models.

Furthermore, consistent with previous observations, the protein chains were found to be more extended in oligomeric states compared to their monomeric counterparts across all systems (see Figure S6). These results underscore the robustness of the LLPS-like behavior across a range of protein concentrations, reinforcing the reliability of our findings.

### Observation of Reentrant phase behavior of Aβ40 at low salt concentration regime

Having established the precedence of LLPS-like phase behavior across a broad spectrum of protein concentrations, our study delved into the potential modulation of microenvironment-induced properties in Aβ40 protein at a concentration of 750 μM. We systematically varied salt concentrations from 25 to 1000 mM under consistent temperature conditions (300 K) to elucidate the intricate relationship between salt concentration and the aggregation of Aβ40 by analyzing the fraction of protein monomers (inversely proportional to the extent of aggregation) which is estimated by calculating the ratio of number of monomers at an instant/total number of protein chains in the system, as depicted in Figure 2A. Intriguingly, we observed a non-monotonic phase behavior within the lower salt concentration range (25-100 mM), contrasting with a more regular trend in the higher salt concentration range (100-1000 mM). Within the lower salt concentration regime, we identified a notable pattern of reentrant phase behavior: initially, the fraction of monomers decreased from 25 mM to 50 mM NaCl salt solution, only to increase upon further elevation to 100 mM, as illustrated in Figure 2B. Conversely, as salt concentration surpassed 100 mM, extending up to 1000 mM, we consistently noted a decline in Aβ40 aggregation propensity, evidenced by a gradual increase in the fraction of monomers, thus following a monotonic trend. These findings underscore the nuanced interplay between salt concentration and Aβ40 phase behavior, highlighting the significance of environmental factors in protein aggregation dynamics, with a maximization of aggregation propensity in presence of 50 mM salt.

**Figure 2:**
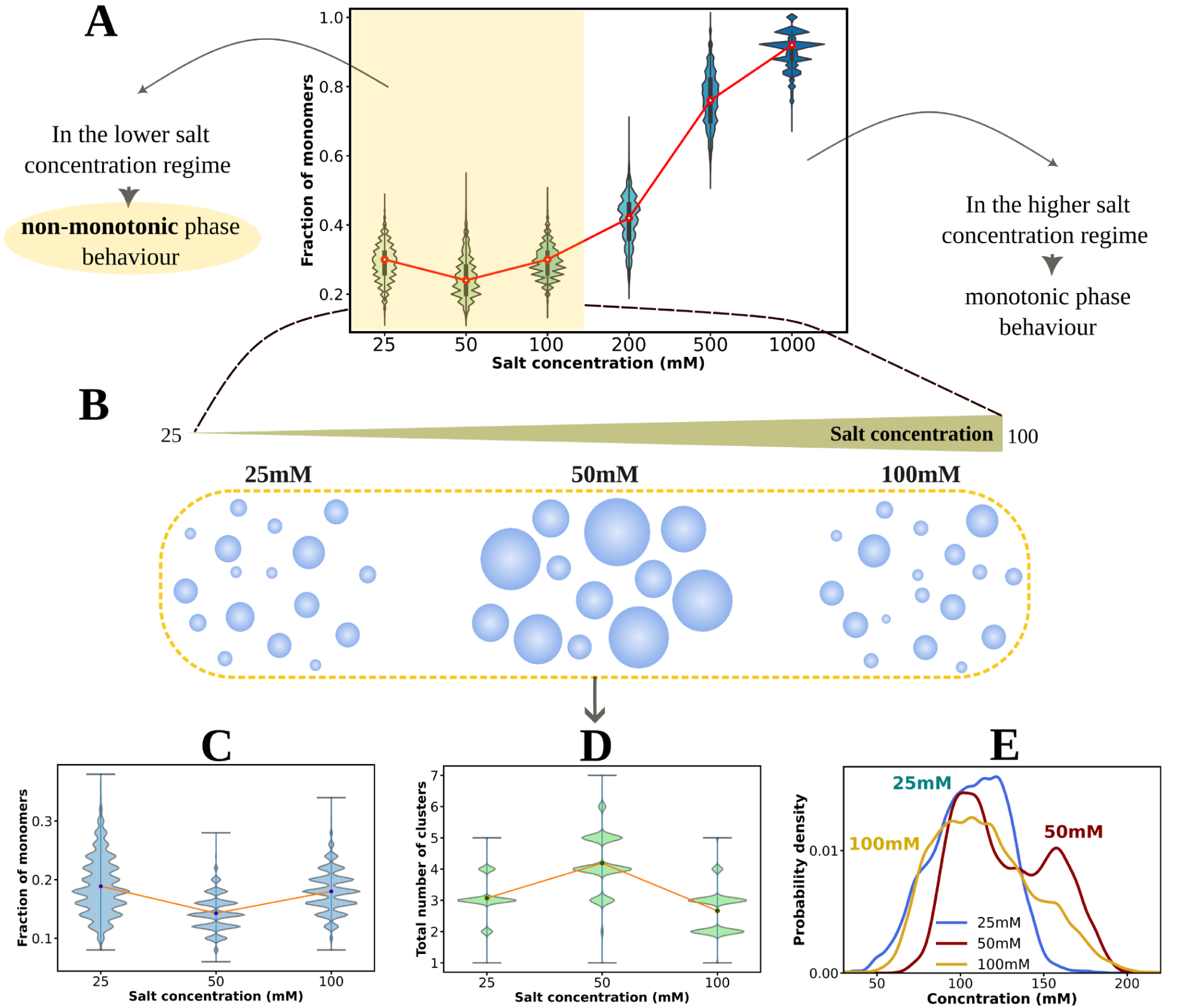
Reentrant condensation of Aβ40 driven by ionic strength variation. A. Probability distribution of fraction of number of monomers (number of monomers at an instant/total number of protein chains in the system) of Aβ40 protein (750 μM) in violin plot with the alteration of salt concentration of the solution from 25 to 1000 mM (25, 50, 100, 200, 500, 1000 mM) at 300 K temperature. B. A schematic representation of Aβ40 reentrant condensation in the lower salt concentration regime (25 to 100 mM). Figure C, D and E. shows Probability distribution of fraction of monomers, total number of clusters (number of protein chains ≥ 6) and protein concentration in dense phase (involving protein hexamer or above) respectively for Aβ40 protein (750 μM at 300 K) in lower salt concentration regime (25, 50 and 100 mM) calculated over the extended trajectories (see method).

To further ascertain the observed non-monotonicity in Aβ40 protein LLPS-like properties, we conducted extensive computer simulations on systems with salt concentrations ranging from 25 to 100 mM. By extending simulation durations to at least 5 μs in each replica, we obtained trajectory data to calculate the fraction of monomers, as depicted in Figure 2C. The trend observed in the extended trajectory reaffirms the non-monotonic nature of salt-induced aggregation propensity, corroborated by the calculation of the total number of aggregates containing six or more protein monomers (Figure 2D).

Moreover, the estimation of protein concentration corresponding to the dense phase (and dilute phase, see Figure S7) revealed intriguing insights. Specifically, we observed an increasing protein concentration in the dense phase from 25 mM to 50 mM salt concentration, indicating a higher tendency for protein aggregation within this range. Conversely, protein concentration in the dense phase decreased with further increases in salt content, particularly above 50 mM to 100 mM, suggesting a reduced probability of protein self-assembly and phase separation within this salt concentration regime. Thus, in the lower regime of ionic strength, Aβ40 exhibits a behavior of reentrant condensation involving an initial transition from lower to higher phase separation tendency, followed by a subsequent reduction.

### Thermodynamic and Kinetic Hallmark of Salt-induced Reentrant condensation of Aβ40

For a more quantitative assessment of the non-monotonic phase separation properties of Aβ40, we utilized the concept of aggregation propensity, as proposed by Tang et al. ^37^. This parameter is typically characterized by two key components: the degree of clustering and the degree of collapse. The degree of clustering, as defined by equation 4, quantifies the ratio of the average number of protein chains in the dense phase (*N_dense*) to the total number of proteins *(N_total*) in the simulation box. On the other hand, the degree of collapse, described by equation 5, is determined by the ratio of the average solvent-accessible surface area (SASA) of all protein chains in the initial state (*SASA_inital*) to that of the final configuration (*SASA_final*). These parameters offer a comprehensive means to quantitatively evaluate the propensity of Aβ40 protein to aggregate and undergo phase separation under varying salt concentrations.

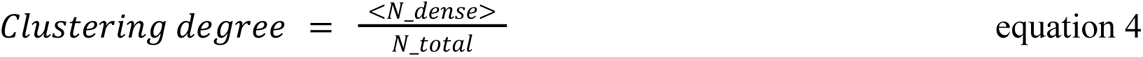

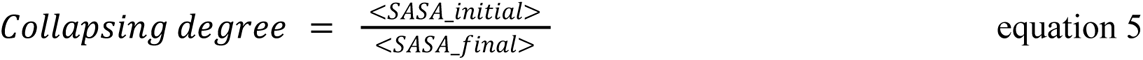

The higher propensity of protein aggregation is indeed reflected by elevated values of both parameters, the clustering degree (indicating more protein chains in the dense phase) and the collapsing degree (suggesting reduced SASA of the final configuration due to increased condensation). As illustrated in Figure 3A, the probability distribution of the clustering degree of Aβ40 protein captures the non-monotonic trend in response to alterations in solution ionic strength from 25 to 100 mM. Notably, at a medium of 50 mM NaCl, the protein exhibits increased susceptibility to aggregation. However, further increases or decreases in salt concentration diminish its phase separation propensity. This observation underscores the intricate and nonlinear nature of Aβ40 phase behavior within the 25-100 mM salt regime.

**Figure 3:**
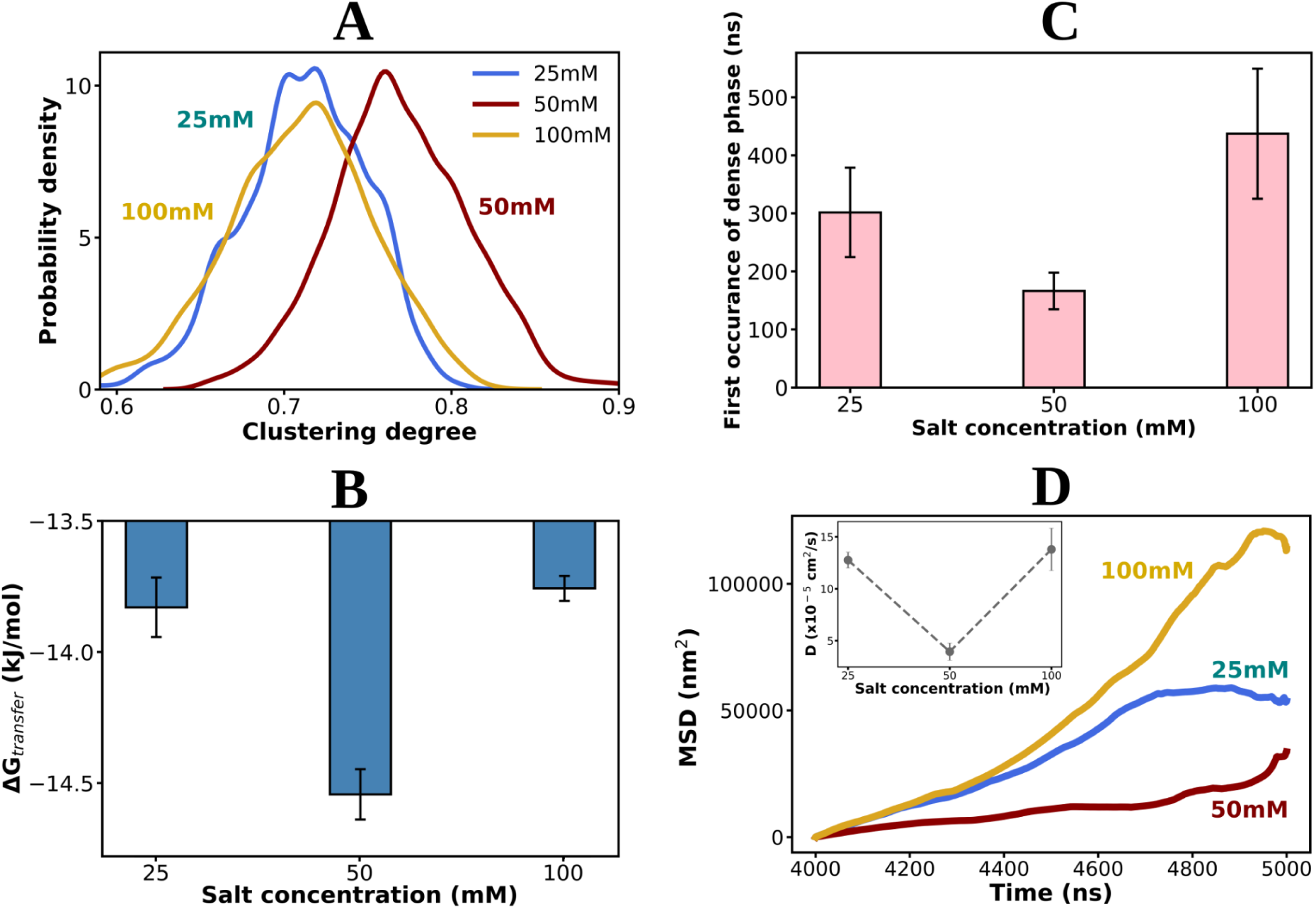
Effect of altering ionic strength in the thermodynamics and kinetics related to the phenomenon of reentrant condensation. A. Probability distribution of clustering degree (see text, equation 4) for Aβ40 protein (750 μM) with the change of NaCl salt concentration 25, 50 and 100 mM at 300 K. Figure B and C represent the change in transfer free energy (ΔGtrans, see text, equation 6) and the time of first occurrence of dense phase respectively of the protein (750 μM at 300 K) in three aqueous saline media 25, 50 and 100 mM in bar plot representation. The errors are calculated by averaging over replicas. Figure D. shows the time profile of mean square displacement of protein corresponding to 25, 50 and 100 mM salt concentrations (for one of the three replicas in each case). Inset shows the change of diffusion coefficient (D) for all the three saline solutions of Aβ40 (750 μM) at 300 K. The errors are calculated by averaging over replicas.

Moreover, the estimation of the degree of collapse, as depicted in Figure S8, further corroborates the reentrant phase behavior of Aβ40 in response to varying salt concentrations within the 25-100 mM range. Together, these findings provide compelling evidence for the nuanced and complex interplay between salt concentration and Aβ40 aggregation propensity, shedding light on the underlying mechanisms governing its phase separation dynamics. Such insights are crucial for understanding the pathological processes associated with protein aggregation, under the exposure of drastic change of environmental conditions.

We further explored the thermodynamic aspect of the nonlinear mode of protein oligomerization propensity, under the influence of variable ionic strength. To achieve this, we calculated the excess transfer free energy (ΔG_trans_) required for transition a protein chain from the dilute phase to the dense droplet phase, as defined by equation 6^5^ (where R represents the universal gas constant, T denotes the temperature, c_dilute and c_dense refers to the concentration of protein in dilute and dense phases respectively).

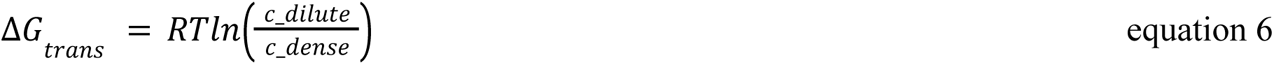

As illustrated in Figure 3B, ΔG_trans_ exhibits its most negative value in the presence of 50 mM salt solution, compared to solutions with 25 and 100 mM salt concentrations. This indicates that the formation of a condensate is most favorable in the system containing 50 mM salt concentration, as the transfer of a protein chain from the dilute to the dense phase becomes most facile there. Hence, the variation in ionic strength significantly influences the thermodynamics of chain exchange between the two phases, thereby crucially regulating the phenomenon of liquid-liquid phase separation (LLPS). Together this observation underscores the pivotal role of ionic strength in modulating the thermodynamic landscape of protein phase behavior, providing valuable insights into the underlying mechanisms governing LLPS dynamics.

Our study unveils the multifaceted influence of changing salt concentration on both the thermodynamics and kinetics of phase separation and aggregation propensity in Aβ40 protein. Figure 3C illustrates the onset-time of first occurrence of the dense phase across the range of 25 to 100 mM NaCl salt solutions containing 750 μM Aβ40 protein at a temperature of 300 K. Intriguingly, a remarkably early onset of condensate formation is observed at 50 mM NaCl, gradually delaying as the ionic strength varies either from 50 mM to 25 or from 50 mM to 100 mM. To further elucidate the dynamics of protein chains over time under these different saline conditions, we characterized the mean square displacement^38^ (MSD) of the proteins, as shown in Figure 3D. It is evident that proteins in the 50 mM NaCl medium exhibit the slowest molecular diffusion compared to the other saline systems, namely 25 or 100 mM. The calculation of diffusion coefficient (D) indicates a decrease in its value upon transitioning from 25 mM to 50 mM salt concentration, followed by an increase when the salt concentration is further increased from 50 to 100 mM (Figure 3D inset). The diminished diffusion of protein chains in a 50 mM saline solution suggests heightened involvement of proteins in self-assembly, consequently slowing down the corresponding dynamics. Collectively, these findings underscore the phenomenon of reentrant condensation exhibited by Aβ40 protein within the low salt concentration regime, influenced by both thermodynamic and kinetic factors.

Very interestingly, a closer observation of previously reported experimental data (figure 1B of the article by Wang et al. ^39^) suggests an occurrence of a similar non-monotonic trend of Aβ40 fibrillation propensity in the similar range of salt concentration, although this aspect has not been attended to in that investigation. In particular, via inspection of the data provided in that study, for the solution of Aβ40 at pH 7.1, it is evident that on the alteration of NaCl salt concentration from 0 mM to 42.9 mM, the t_1/2_ values of the fibrillation kinetics gradually decreased first but upon further rise of salt concentration beyond 42.9 till 71.4 mM the t_1/2_ values again increased indicating lesser propensity of Aβ40 aggregation. Similar trend was also indicated by Krainer et al. showing significant phase separation at the low salt concentration of 50 mM while dissolution of condensates at the relatively higher salt concentration (above 500 mM) for a wide range of proteins FUS, FUS G156E, TDP-43, Brd4, Sox2 and A11 at pH ∼7^23^. Reentrant phase behavior, characterized by a transition from low to high phase separation tendency followed by a subsequent reduction, has also been previously reported in various systems including colloid, DNA, RNA, and certain proteins due to modulation of solution ionic strength^20,22^. Proteins such as FUS, TDP-43, Brd4, Sox2, and Annexin A11 have been observed to undergo condensation at low salt concentrations (50 mM), dissolution at intermediate concentration (above 500 mM) and transition back into a phase-separated state at high salt concentrations (beyond 1.5 M)^23^. Conversely, for colloids, nucleic acids, and some globular proteins (e.g., BSA, HSA, OVA, and β-LG), reentrant condensation behavior has been reported, involving resolubilization upon the introduction of additional salts following phase separation^20,22^. Some studies have hypothesized the formation of biomolecule-salt overcharged complexes as the key reason for reentrant behavior, while others attribute it to the stabilization of non-ionic and hydrophobic interactions^20,22,23^. In this context, the observations presented in the current setup highlight the complex interplay between biomolecular interactions and environmental factors, underscoring the need for comprehensive studies to elucidate the underlying mechanisms governing phase behavior in biological systems.

### Altered protein electrostatics drives the salt-concentration dependent reentrant phase behavior of Aβ40

Achieving a comprehensive understanding of the non-monotonic phase behavior of Aβ40 protein in altered saline medium requires delving into the specific influence of salt-screening on the protein. Conventional dielectric-continuum based models are insufficient due to the intricate charge distribution and geometries involved^20^. While alternative theoretical frameworks of Debye-Hückel and its derivatives address complex geometries, they fall short in scenarios with significant screening charges where ion-ion correlations become critical^20^. Additionally, the linearized Poisson-Boltzmann theory underestimates the influence of salt, especially in the case of monovalent ions^22^. Therefore, more fine grained molecular interpretations are essential for a deeper understanding^20^.

In our investigation (depicted in Figure 4A), we focused on the origin of non-monotonic phase behavior by examining the specific influence of salt-screening on the protein. Recent studies have emphasized the altered protein electrostatics due to the screening effect of salt, which leads to abnormal protein-protein aggregation. The strong screening effect of salt partially neutralizes the surface charge of the molecule, reducing surface repulsion and driving aggregation^40,41^.

**Figure 4:**
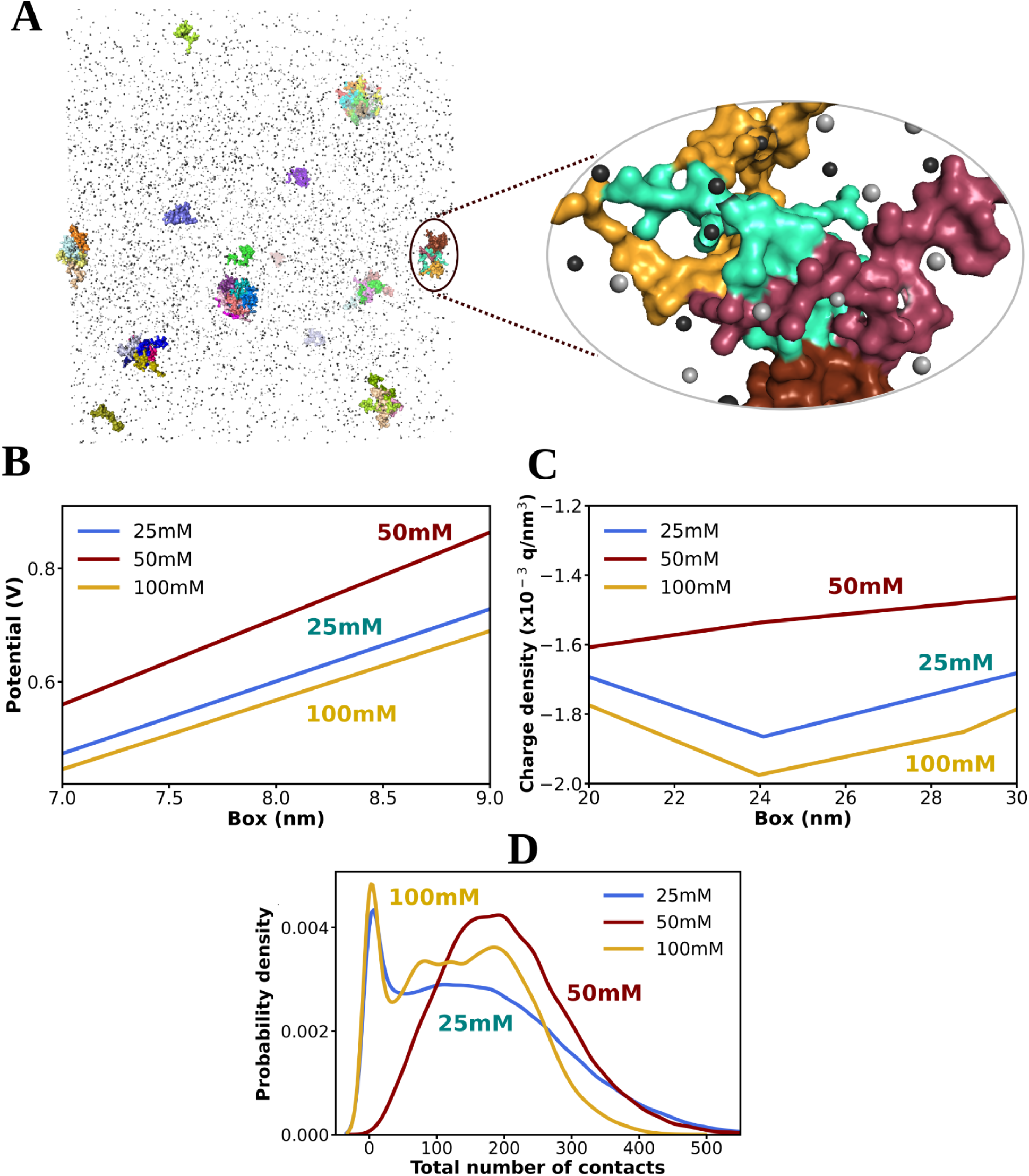
Altered protein electrostatics drives salt-induced reentrant condensation of Aβ40. A. Schematic representation of Aβ40 protein forming oligomers in presence of NaCl salt species. Proteins are shown in surface representation by coloring by chains. Salt cations and anions are shown in dark silver and light silver coloration. In the right hand side the zoomed in view is being represented. Figure B. and C. represent the variation of potential and charge of protein across the box (see method) for three salt concentrations 25, 50 and 100 mM at 300 K temperature. Figure D. represents the probability distribution of total number of interchain contacts (within the cutoff of 0.8 nm) of the proteins (in oligomers) for the alteration of salt concentration from 25 to 50 to 100 mM at 300 K temperature.

To explore this phenomenon, we analyzed the alteration of protein electrostatics with varying salt concentration. The mode of electrostatics of protein molecules in different environments were derived by calculating electrostatic potential and charge of protein across the box (see method). Figure 4B illustrates the profile of the potential corresponding to the biomolecule across the simulation box, showing non-monotonic changes in protein potential in the 25-100 mM saline regime (see method). On a related note, the overall charge is calculated by summing the charges in regular slices corresponding to the protein (see method). Figure 4C demonstrates the alteration of protein charge profile across the box, which also follows a non-monotonic trend with changes in salt concentration in the range of 25 to 100 mM.

Increasing the salt concentration from 25 to 50 mM decreases the protein charge (becomes less negative) due to higher charge screening by salt, favoring closer approximation of biomolecules and leading to higher aggregation propensity in the 50 mM saline condition facilitated by higher probability of intermolecular interaction (Figure 4D). Conversely, changing the salt concentration from 50 to 100 mM increases the protein charge content, promoting higher intermolecular repulsion and resulting in lesser protein aggregation.

Overall, the non-monotonic alteration of protein electrostatics guided by the interplay of changing ionic strength in the 25-100 mM saline regime give rise to the nonlinear phase behavior of Aβ40 protein, particularly reentrant condensation. Interestingly, this phenomenon has been observed in earlier experimental studies based on electrophoresis measurements for proteins or peptides in the presence of trivalent ions^20,22,25^, but our study demonstrates that reentrant behavior can be guided by monovalent ions that are efficiently seeded from altered protein electrostatics by salt-interplay. This highlights the intricate interplay between protein electrostatics and salt concentration in driving phase behavior, providing valuable insights into the underlying mechanisms of protein aggregation dynamics.

### Salt-mediated Electrostatic Modulation of Aβ40 N-Terminus Controls Hydrophobic Contact and Drives Reentrant Phase Behavior

To comprehend the pivotal interactions governing the phase separation of Aβ40 and their modulation during the protein’s reentrant behavior, we investigated the amino-acid pair interactions within densely packed protein chains. Specifically, we aimed to elucidate the impact of microenvironmental conditions on protein-protein interactions, leading to reentrant phase behavior, by estimating the differential residue-wise interprotein contact map across salt concentrations of 25, 50, and 100 mM.

Differential contact maps were computed by first deriving residue-wise interprotein contact probability maps corresponding to specific salt concentrations and then subtracting the contact probability values between different salt concentrations. Notably, Figure 5A illustrates the difference in residue-wise interprotein contact maps between 25 mM and 50 mM salt conditions, revealing an intensified interprotein interaction upon increasing salt concentration. This intensification, indicated by red-colored regions in Figure 5A, primarily involves enhanced interactions among the protein’s central hydrophobic core (CHC) regions (CHC-CHC contact) and between the CHC and secondary hydrophobic regions (SHR) of the proteins (CHC-SHR contact).

**Figure 5:**
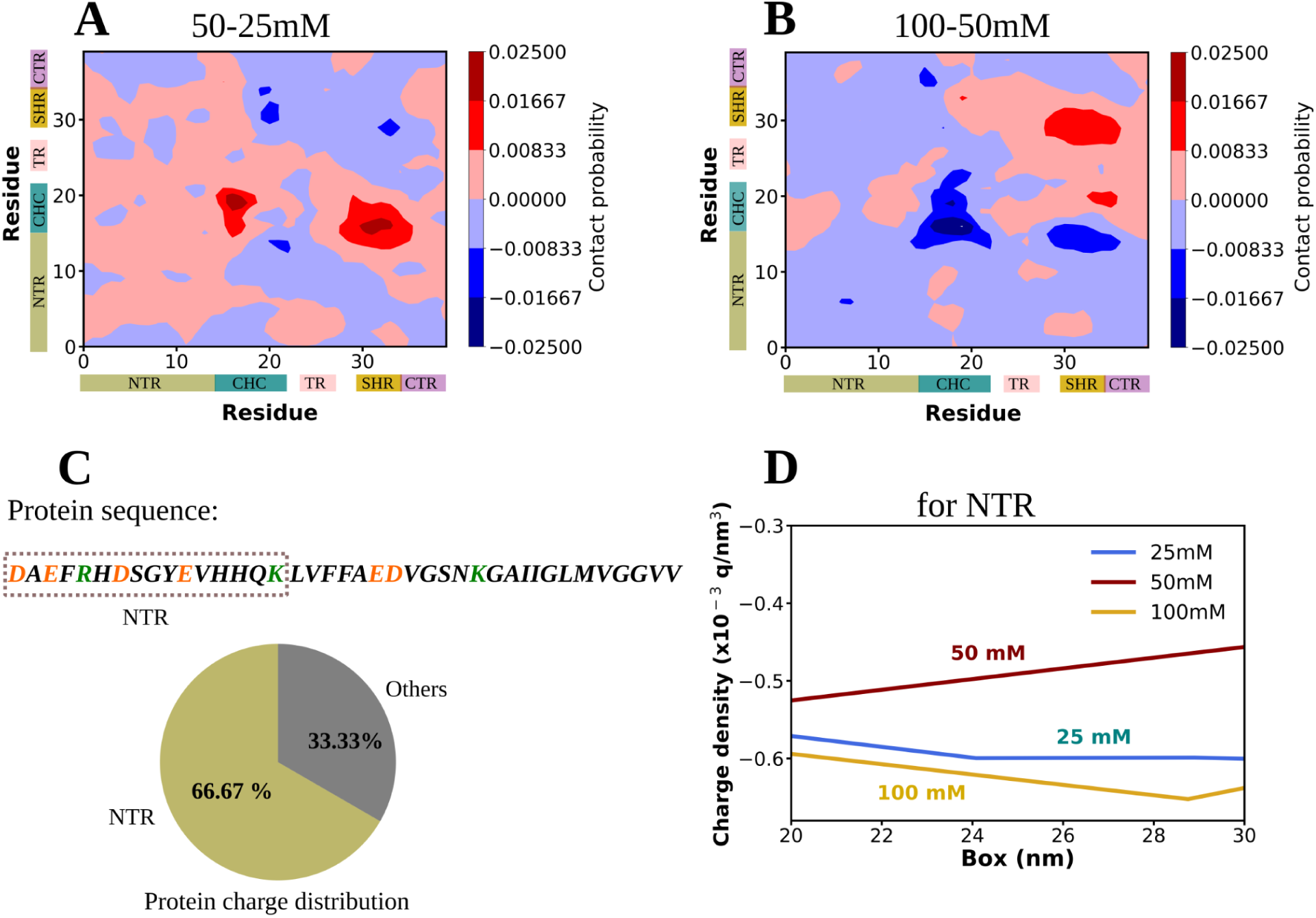
Molecular insights for salt dependent reentrant behavior of Aβ40. Figure A and B shows the differential residue-wise contact maps among the Aβ40 (750μM) protein chains (interchain) in dense phase corresponding to the change in salt concentration from A. 25 to 50 mM (interchain contact probability in 25 mM salt is subtracted from that of in 50 mM salt concentration) and B. 50 to 100 mM (interchain contact probability in 50 mM salt is subtracted from that of 100 mM salt concentration) at 300 K temperature. The different regions of protein is represented as the N-terminal region (NTR, residues 1-16) in khaki, the central hydrophobic core (CHC, residues 17-21) in teal, the turn region (TR, residues 24-27) in pink, the secondary hydrophobic region (SHR, residues 30-35) in golden, and the C-terminal region (CTR, residues 36-40) in purple color. Figure C. shows the entire Aβ40 protein sequence where the acidic and basic (charged residues) residues are colored with orange and green colors respectively. The NTR part which contains most of the charged residues, is highlighted with a dashed lined box. The pie chart depicts overall charged residue distribution in NTR, (66.67%) and the other parts (33.33%) of the protein. Figure D represents the profile of charge density corresponding to NTR of the Aβ40 protein (750μM) in varied concentrations of aqueous NaCl solution (25, 50 and 100 mM) at 300 K temperature.

Prior studies have emphasized the crucial role of CHC-CHC and CHC-SHR contacts in Aβ40 oligomerization^21,42^. The observed increase in these interactions with salt concentration variation from 25 to 50 mM suggests that the heightened propensity for protein aggregation at 50 mM NaCl is stabilized by favorable hydrophobic contacts. Conversely, altering salt concentration from 50 to 100 mM (Figure 5B) diminishes the contact probabilities associated with key protein interactions (reflected by the blue regions in Figure 5B) responsible for aggregation, indicating a reentrant phase behavior.

Further exploration into the driving forces behind the variation in the relative stability of protein-protein hydrophobic interactions across salt concentrations led us to investigate salt-concentration dependent alterations in protein electrostatics. We focused on the N-terminal region (NTR), which contains a significant portion of charged residues (Figure 5C). Our analysis revealed a decrease in the charge of the NTR region from 25 to 50 mM NaCl, while a subsequent increase occurred from 50 to 100 mM NaCl (Figure 5D).

At 50 mM NaCl, the charge of the NTR region is largely screened, resulting in reduced protein repulsion (facilitating closer approach of the protein molecules) and stabilized intermolecular contacts involving favorable hydrophobic interactions with CHC and SHR regions. Consequently, salt-dependent alterations in electrostatics of the NTR region non-monotonically stabilize interprotein hydrophobic interactions, leading to reentrant condensation of Aβ40.

### Reentrant condensation of Aβ40 recurs in temperature space

Building upon the crucial effect of microenvironmental conditions, particularly alterations in ionic strength on Aβ40 phase behavior, we extended our investigation to explore the influence of solution temperature on the aggregation propensity of Aβ40 protein. We simulated multi-chain behavior of the amyloid protein (750 μM protein in 50 mM NaCl salt solution) for 2.5 μs in three replicas, varying temperatures from 270 to 330 K at regular intervals of 10 K (Figure S9). Intriguingly, similar to the case of salt, a phenomenon of reentrant condensation was observed for Aβ40 protein in response to temperature variation. Figure 6A illustrates the change in fraction of monomers with temperature alteration, revealing a non-monotonic trend where the fraction initially decreases from 270 K to 280 K, then increases with further temperature rise. The same trend was retained when extending simulations to at least 5 μs for temperatures ranging from 270 to 290 K (270, 280, and 290 K), as shown in Figure 6B. Furthermore, analysis of the fraction of monomers for the extended trajectories (Figure 6B) and estimation of total number of clusters (Figure S10) supported the observed reentrant phase behavior of Aβ40 with temperature in the range of 270-290 K. Protein concentration in the dense phase (Figure 6C) showed an increase (and dilute phase (in Figure S11) showed decrease) at 280 K but decreased (dilute phase (in Figure S11) increased) at 290 K, indicating a temperature-dependent modulation of aggregation propensity following a non-monotonic trend in the low temperature regime of 270-290 K.

**Figure 6:**
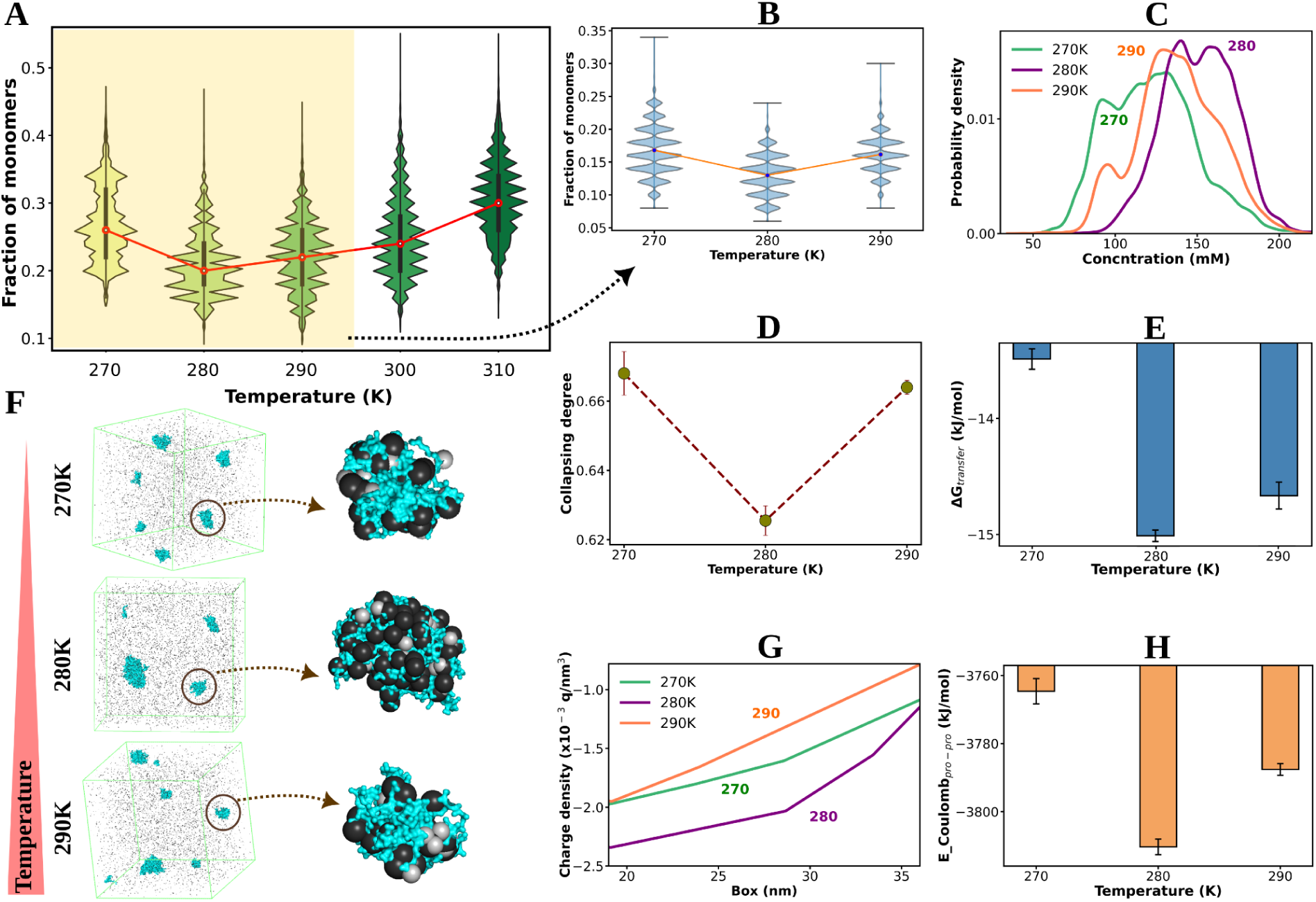
Temperature induced reentrant condensation of Aβ40. A. Probability distribution of fraction of number of monomers (number of monomers at an instant/total number of protein chains in the system) of Aβ40 protein (750 μM) dissolved in 50 mM aqueous NaCl solution are shown in violin plot with the alteration of temperature from 270 to 310 K with the interval of 10 K (i.e. 270, 280, 290, 300 and 310 K). Figure B, C, D and E shows Probability distribution of fraction of monomers, protein concentration in dense phase, the change in collapsing degree and transfer free energy (ΔG_trans_) respectively for Aβ40 protein (750 μM) in 50 mM salt concentration for 270-290 K temperature range (270, 280 and 290 K) calculated over the extended trajectories (see methods). Figure F shows representative simulation snapshots for the systems containing 750 μM Aβ40 protein in 50 mM NaCl solution for 270, 280 and 290 K temperature (top to down). Proteins are shown in cyan colored surface representation and the salt ions (cation: dark silver, anion: light silver) are shown in spheres. In the right hand size a zoomed in view of an protein oligomer surrounded by salt ions are being represented. Figure G. shows the alteration of protein charge across the simulation box for the similar system on varying the temperature from 270 to 280 to 290 K (calculated over longer trajectories). Figure H. represents the estimation of Coulombic interaction energy among proteins (750 μM Aβ40 in 50 mM NaCl salt) with the change in temperature in the range of 270 to 280 K (calculated over longer trajectories). The error bars in figure D, E and H are shown in black colored vertical lines estimated by averaging over different replicas.

The degree of collapse (Figure 6D) and clustering degree (Figure S12) further affirmed the thermoresponsive reentrant phase behavior of Aβ40, guided by the modulation of thermodynamic properties associated with Aβ40 aggregation (Figure 6E). Specifically, the free energy of transfer (ΔGtrans) decreased considerably from 270 to 280 K, making the process of condensation more favorable. Conversely, increasing the temperature from 280 to 290 K led to a less negative ΔGtrans, resulting in a lesser extent of aggregation.

Interestingly, our mechanistic insight revealed that the alteration in salt interaction with protein with temperature variation modulated the protein charge via screening effect. Figure 6F (and Figure S13) shows an increase in accumulation of salt species near the protein surface at 280 K compared to 270 or 290 K. Therefore the charge of protein decreased from 270 to 280 K due to higher screening effect (Figure 6G), favoring more protein-protein interaction and enhancing aggregation propensity at 280 K (Figure S14 represents increased interprotein contacts). Conversely, a further increase in temperature from 280 to 290 K led to an increase in protein charge, decreasing the propensity of protein condensation and resulting in the reentrant phenomenon. Figure 6H depicts the impact of temperature on Coulombic interaction of protein due to temperature-induced protein electrostatics modulation, corroborating the observed non-monotonic changes in the total number of interprotein contacts with temperature (Figure S14).

The highest propensity of Aβ40 condensation at around 280 K as suggested by reentrant condensation in our investigation, has been previously noted in former report^43^ indicating the occurrence of relatively stable soluble Aβ oligomers at the low temperature of around ∼4°C i.e. 277 K at low ionic strength and pH ∼7, a condition akin to the present article. A previous report by Stine et al. in 1998 has also highlighted temperature of ∼4°C as an optimum temperature of amyloid oligomer formations^44^. In this study the authors have very clearly depicted through atomic force microscopy (AFM) the well formed oligomers of amyloid beta protein after incubation of protein solution at 4°C for 24 hrs while fibril formation accelerated at 37°C. In this line a former inspection by Ahmed et al. also suggests that low temperature like 4°C can stabilize Aβ oligomers which are largely more toxic in nature to neurons compared to the other forms like protofibrils and fibrils^45^. On temperature rise the stable oligomers can convert to protofibrils and fibrils eventually. Together these support our investigation involving enhanced tendency of Aβ40 protein to form condensates favorably at the temperature around 280 K.

Similar to the scenario of salt driven reentrant Aβ40 condensation, during the alteration of temperature the non-monotonic trend of protein-protein hydrophobic interactions involving key aggregation causing protein regions CHC and SHR has been observed (Figure S15 A-B). From 270 to 280 K more intense CHC-CHC and CHC-SHR interprotein interaction has been observed (the red regions of the differential contact map in Figure S15 A) while further increase of temperature from 280 to 290 K decreases the strength of interprotein interaction (the blue regions of the differential contact map in Figure S15 B) previously stabilized through hydrophobic contacts. It is due to non-monotonic alteration of electrostatics of the most charged NTR part of the protein (Figure S15 C), the variability in the propensity of interprotein interactions is originated. Change of temperature from 270 to 280 K decreases the charge of Aβ40 NTR which can facilitate closer approximation of the protein chains due to reduction of interprotein repulsion leading to favorable hydrophobic contacts crucial for Aβ40 condensation. So, the altered protein electrostatic interactions driven by temperature changes (in the range of 270 to 290 K) guide the thermoresponsive reentrant Aβ40 condensation stabilized by hydrophobic interprotein interactions following a non-monotonic trend.

## Conclusion

The present investigation underscores the critical role of surrounding microenvironments in shaping the liquid-liquid phase separation (LLPS) properties of the amyloid protein Aβ40, implicated in Alzheimer’s disease (AD). Our study highlights the context-dependent nature of LLPS properties, wherein microenvironmental factors exert significant influence on the aggregation behavior of Aβ40 beyond its chemical framework. Interestingly, our findings reveal that modulation of salt concentration and temperature induces reentrant behavior in condensation properties, driven by altered intermolecular interactions. At a molecular level, the charged residues at N-terminus show non-monotonic screening by the salt-concentration, which modulates the hydrophobic contacts in Aβ40 and drives reentrance in phase behavior. This sheds light on the early stages of condensation of pathogenic Aβ40 protein in response to variable environmental states, alongside the thermodynamic and kinetic properties of condensates. Understanding the specific LLPS behavior of Aβ40 expands the scope of its druggability and holds promise for addressing challenges in the field of health and disease.

Experimental studies of Aβ condensation are inherently challenging due to the protein’s high structural fluctuation and rapid aggregation process. Moreover, modulation of amyloid aggregation by various microenvironmental conditions adds complexity to the condensation process. While previous studies have explored the molecular nature of amyloid aggregation, the precise effects of different microenvironmental scenarios, known to strongly influence macroscopic aggregation properties, remain unclear. In this respect, our findings emphasize the importance of specific microenvironmental conditions encountered by biomolecular condensates. We observe a non-monotonic trend in Aβ40 aggregation propensity with variations in salt concentration and temperature, guided by altered intermolecular interactions under specific solution conditions. Electrostatic interactions, guided by these conditions, play a pivotal role in shaping non-linear protein-protein interactions through altered charge compensation, ultimately influencing Aβ40 phase behavior by non-monotonically stabilizing the interprotein hydrophobic interactions. Our research highlights the potential to adjust protein interactions by compensating and overcompensating protein charges through fine-tuning environmental conditions. Understanding the effects of specific solution conditions on phase separation processes is crucial for predicting, rationalizing, or modulating protein characteristics in designing therapies against phase separation-related pathologies. Our findings illuminate the importance of solution condition dependent modified protein electrostatics influencing protein hydrophobic interactions as fundamental drivers in the condensation process, with significant implications for abnormal functionality, drug targeting, and material characteristics of biomolecular condensates.

### Simulation Model and Methods

Major computer simulations in this article were performed with Aβ40 protein using Martini version3 coarse-grained model^27^ utilizing Gromacs software of version 20xx^46^. Since the regular Martini 3 parameters of the coarse-grained model are mostly developed for single-domain and multi-domain proteins, these unchanged parameters are known to underestimate the chain size of such systems and to overestimate protein-protein interactions. Hence we first optimized the Martini3 parameters (corresponding to water-protein interactions) both at the monomeric and dimeric protein level based on the results obtained from the corresponding atomistic simulations reported in previous studies (atomistic monomeric data are obtained from D. E. Shaw research group^47^ and atomistic dimeric data are collected from former study by Sarkar et al.^21^).

### Refining the Martini 3 coarse-grained model tailored to Aβ40 protein via Benchmarking against atomistic simulation

Previous efforts to simulate IDPs involved adjusting the Martini-3 force field by fine-tuning water-protein interactions, specifically the σ and ε parameters of Lennard-Jones interactions^5,48,49^. In this study, we adopt a similar approach involving refinement of the ɛ component of the water-protein Lennard-Jones interactions based on atomistic simulation with monomer and dimer of Aβ40 protein with Charmm36m force field to better model the IDP (equation 4).

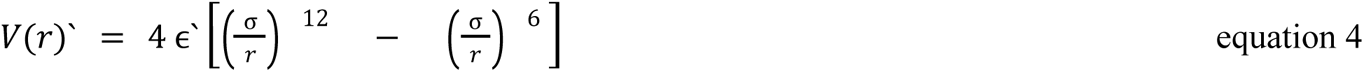

Where ɛ′ = λɛ and λ represents the scaling parameter requiring optimization. Modifying ɛ adjusts the potency of the water-residue interactions.

Initially we conducted coarse-grained simulations of single-chain Aβ40 immersed in water within the simulation box^21^ of dimension 8.3×8.3×5.9 nm^3^ maintaining NaCl salt concentration at 50 mM^47^ while adding few additional ions within the system to maintain electroneutrality. The protein,water and salt were modeled using Martini-3 resolution. To achieve the optimal scaling parameter for water-protein interactions within Martini-3, particularly tailored to Aβ40, we conduct coarse-grained simulations employing two Aβ40 chains with varying λ values. Each of the simulation is performed in three replicas for at least 1 μs long in each of the cases (time step = 0.02 ps) and analysis is performed over the trajectories of all the three replicas corresponding to each of the λ values. Through these simulations, we derive the average R_g_ values, aiming to match those obtained from the atomistic simulation trajectory of Aβ40 monomer obtained from D. E. Shaw research group^47^. The atomistic simulation is performed using CHARMM36m force field for 30 μS long in 50 mM aqueous NaCl solution^47^. Achieving this alignment involves fine-tuning λ, as depicted in Figure S1.

To further verify the suitable value of λ in modeling protein-protein interactions, we conduct simulations of the dimerization process of Aβ40 with varying λ values using Martini 3 force field. We commence by positioning two chains, devoid of any enforced secondary structure, randomly within the simulation box (of dimension 8.3×8.3×5.9 nm^3^). We maintained an interchain distance more than 0.8 nm^21,24^ which we utilized as the cutoff distance to classify the chains as bound. By varying the λ parameter a series of simulations are performed in a 50 mM monovalent saline medium making sure the charge neutrality of the system. Each of the simulations corresponding to a particular λ value are carried out for at least 1 μS (time step of 0.02 ps) and is repeated thrice. The analysis for each λ value was performed by combining all the three trajectories together corresponding to a λ. As our focus lies in leveraging multi-chain simulations to investigate the liquid-liquid phase separation (LLPS) of Aβ40 via Martini 3, we employ the duration of two all-atom monomers staying bound to each other as the benchmark. Utilizing this defined cutoff of 0.8 nm, we compute the percentage of bound conformations between the Aβ40 monomers for different λ values within the coarse-grained model. Additionally, we compare these results with data derived from previously reported atomistic simulations^21^ of Aβ40 dimers. The atomistic simulation of Aβ40 dimers are performed in the Charmm36m force field^47^ (for protein and ions) in the aqueous medium containing 50 mM NaCl salt concentration using the Charmm-TIP3P^50^ model of water. Atomistic dimer simulations are performed starting from the randomly placed protein chains kept at a distance apart from each other (1, 2 and 4 nm i.e. more than interaction cut-off of 0.8 nm) in the simulation box of dimension 8.2×8.2×5.7 nm^3^. Total three sets of atomistic dimer simulations are reported in 2 replicas each for at least 500 ns (total 3 μS). The overall trajectory data is utilized for further calculations and compared with the CG simulation results performed here with the two Aβ40 chains modeled with Martini-3 force field parameters by varying λ. As illustrated in Figure S2, we observe a consistent agreement in the percentage of bound conformations between coarse-grained (performed here) and atomistic simulations (reported by Sarkar et al.^21^) across various λ values. Consequently, by evaluating both the radius of gyration and the percentage of bound conformations, we determine λ = 0.995 as the optimal scaling parameter for water-protein interactions in modeling Aβ40 with Martini 3.

### Setup for aggregation Simulations

To investigate the phase behavior of Aβ40 protein in response to the alteration of protein, salt concentration and temperature, a series of simulations have been carried out with multiple Aβ40 protein chains modeled with optimized (explained above) Martini3 parameters utilizing GROMACS software of version 20xx^46^.

#### System details

We simulated the aggregation behavior of multiple copies Aβ40 protein chains randomly dispersed in a cubic box of dimension 48×48×48 nm^3^ by varying the protein concentration from 500 to 750 to 1000 μM concentration in 50 mM aqueous NaCl medium at 300 K temperature. Each system was solvated by water and required number of additional monovalent ions (either Na^+^ or Cl^-^) to maintain the necessary salt concentration as well as overall charge neutrality of the system. As a key investigation, we explored the effect of changing salt concentration in the aggregation behavior of Aβ40 maintained at 750 μM concentration at 300 K temperature. In particular, we varied the NaCl concentration from 25 to 1000 mM (25, 50, 100, 200, 500 and 1000 mM). Next we have also inspected the influence of altering solution temperature for the system of 750 μM Aβ40 dissolved in 50 mM aqueous NaCl salt solution. The specifics of the simulation setup are provided in Table 1. The temperatures are varied in the range of 270 to 330 K with a regular interval of 10 K i.e. 270, 280, 290, 300, 310, 320 and 330 K respectively. For all the cases box size are kept constant as 48×48×48 nm^3^.

#### Simulation methods

We started our simulation with randomly dispersed protein chains modeled with the scaled Martini 3 parameters, dissolved in the aqueous salt solution at desired concentration and at particular temperature. We commence with energy minimization employing steepest gradient descent until reaching a tolerance of 10 kJ mol^−1^ nm^−1^. Subsequently, we conduct NVT simulations at specified temperature, employing the v-rescale thermostat for 5 ns with a time step of 0.01 ps. This step is succeeded by NPT simulations at that particular temperature and 1 bar pressure, utilizing a v-rescale thermostat and Berendsen barostat for 5 ns with a time step of 0.02 ps.

Subsequently, we perform coarse-grained molecular dynamics (CG MD) simulations employing a velocity-verlet integrator with a time step of 0.02 ps, utilizing a v-rescale thermostat and a Berendsen barostat to maintain desired temperature and pressure (1 bar) respectively. Both Lennard-Jones and electrostatic interactions are truncated at 1.1 nm. Coulombic interactions are computed using the Reaction-Field algorithm with a relative dielectric constant of 15.

For each of the systems we have performed MD simulations with scaled Martini 3 parameters for at least 2.5 μs in three different replicas for ensuring statistical reproducibility. For analysis, the last 1 μs (1.5 to 2.5 μs simulation data) trajectory of each of the replica simulations are concatenated (overall 3 μs). We have observed reentrant condensation of Aβ40 for the salt concentration regime of 25-100 mM (750 μM protein at 300 K temperature) and also upon alteration of temperature in the range of 270-290 K (750 μM protein in 50 mM NaCl concentration). To ensure the statistical reproducibility of the reentrant behavior captured in our simulations at low (25-100 mM) salt concentration and temperature (270 to 290K), we have extended the simulations corresponding to the non-monotonic salt (25-100 mM) and temperature range (270-290 K) at least up to 5 μs long for each of the three replicas corresponding to each of such system. Analysis is performed on the concatenated simulation trajectories of the final 1 μS (4 to 5 μS) of each of the three replicas corresponding to each of such simulation systems.

The mode of electrostatics of protein molecules in different environments are analyzed by calculating electrostatic potential and charge of protein across the box. This can be calculated by mapping atomic positions, which in turn define the electrostatic potential distribution within a system^51^. The overall charge is calculated by summing the charges in regular slices corresponding to the protein. Thereafter the potential has been calculated by integrating two times of this charge distribution. We have calculated the charge and electrostatic potential for the Aβ40 in different microenvironmental conditions. Here the gromacs tool of gmx potential has been utilized for the electrostatics based calculations. Since it does not consider periodic boundary conditions (pbc) for the calculations we have provided pbc corrected trajectories as our inputs.

## Supplementary Information

Supplementary information contains supplemental figures and tables.

## Acknowledgement

We acknowledge support of the Department of Atomic Energy, Government of India, under Project Identification No. RTI 4007. JM acknowledges Core Research grants provided by the Department of Science and Technology (DST) of India (CRG/2023/001426). We acknowledge Abdul Wasim for sharing certain codes that were useful for analysis

## Notes

### Competing Interest Statement

The authors have declared no competing interest.

