## Supplementary material for "Microenvironment Drives Reentrant Condensation of Aβ40": supplental figures

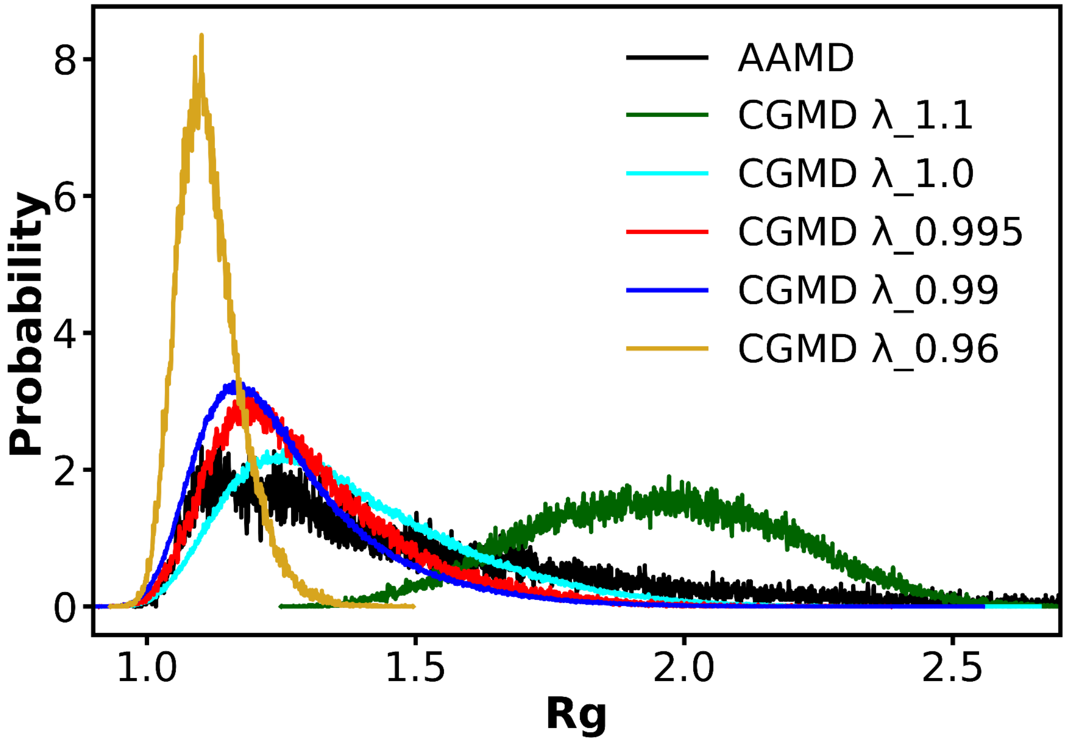


**Figure S1: Scaling of λ value based Aβ40 on monomer simulations.** Figure shows the probability distribution of Aβ40 monomer in 50 mM NaCl solution at 300 K temperature corresponding to the CGMD simulations (utilizing Martini 3 parameters) performed on altering the λ value and atomistic simulation obtained from D. E. Shaw research group[1].

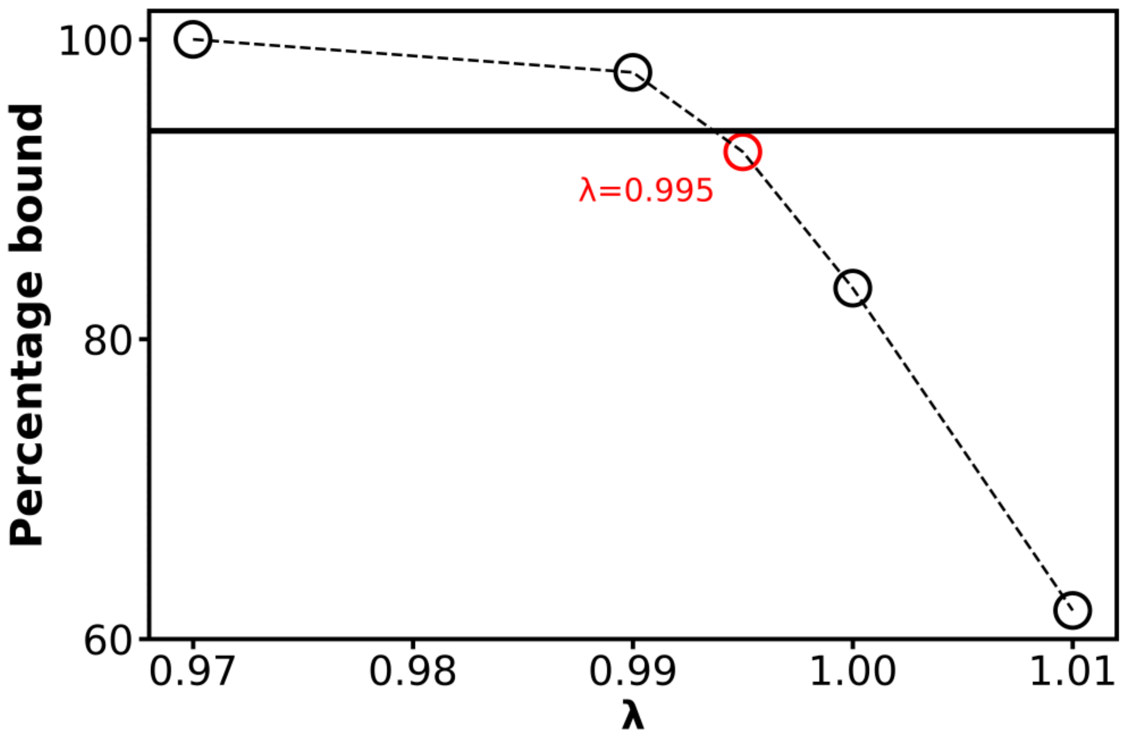


**Figure S2: Scaling of λ value based Aβ40 on dimer simulations.** Figure represents the percentage bound values for the two chains of Aβ40 proteins (see method) in CGMD simulation employing Martini 3 parameters (in 50 mM aqueous solution NaCl salt at 300 K temperature) performed with tuning the values of λ. The horizontal solid line represents the λ value obtained from the atomistic dimer simulations of Aβ40 (50 mM aqueous solution NaCl salt at 300 K temperature) reported previously by Sarkar et al.[2]. The value of λ recapitulates the nearest percentage bound in CGMD simulation with respect to the atomistic simulation has been shown in red colored circle representation.

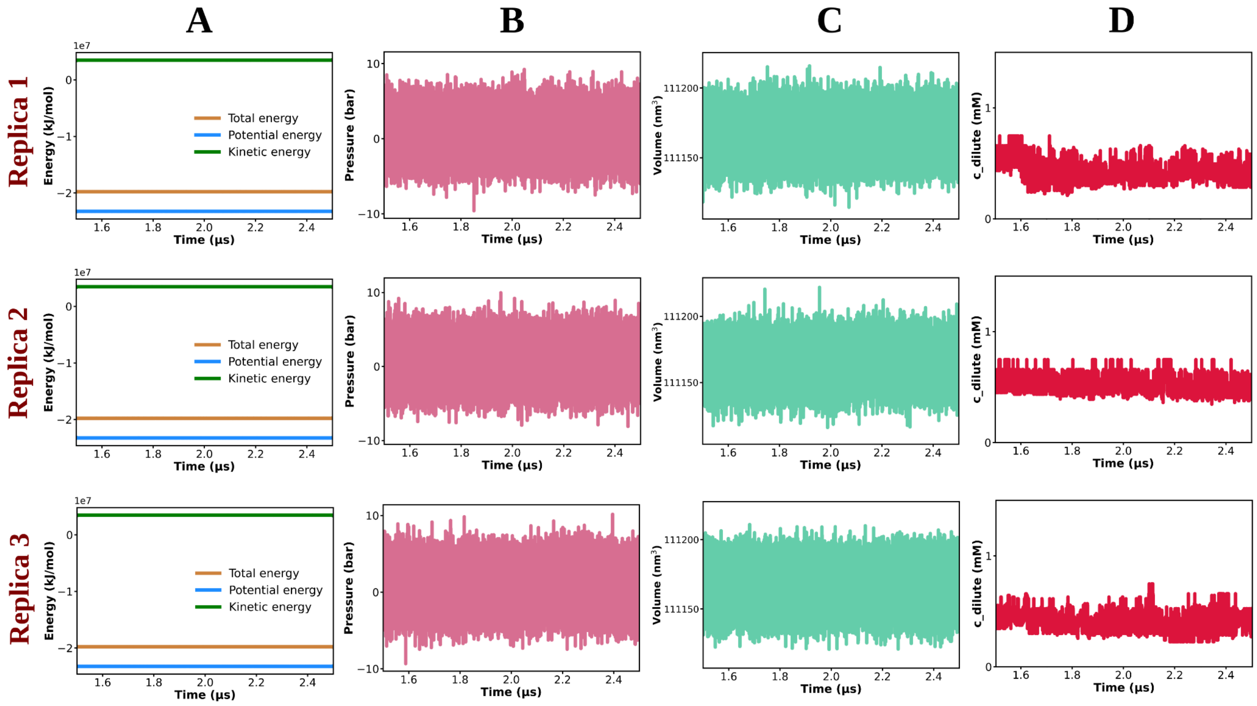


**Figure S3: Systems all well equilibrated during the simulation time.** Figure A. shows the time profile of total energy, potential energy and kinetic energy of the system containing 750 μM of Aβ40 protein in 50 mM aqueous NaCl solution at 300 K temperature corresponding to all the three replicas (shown one by one from top to bottom) for the final 1 μS simulation (1.5 to 2.5 μS) trajectories which are utilized here for analysis. Figure B, C and D represent the time profile of pressure, volume and concentration of protein in dilute phase (c_dilute) for the same system (750 μM protein in 50 mM NaCl solution at a temperature of 300 K) corresponding to final 1 μS trajectory of each of the three replicas (shown top to bottom consecutively).

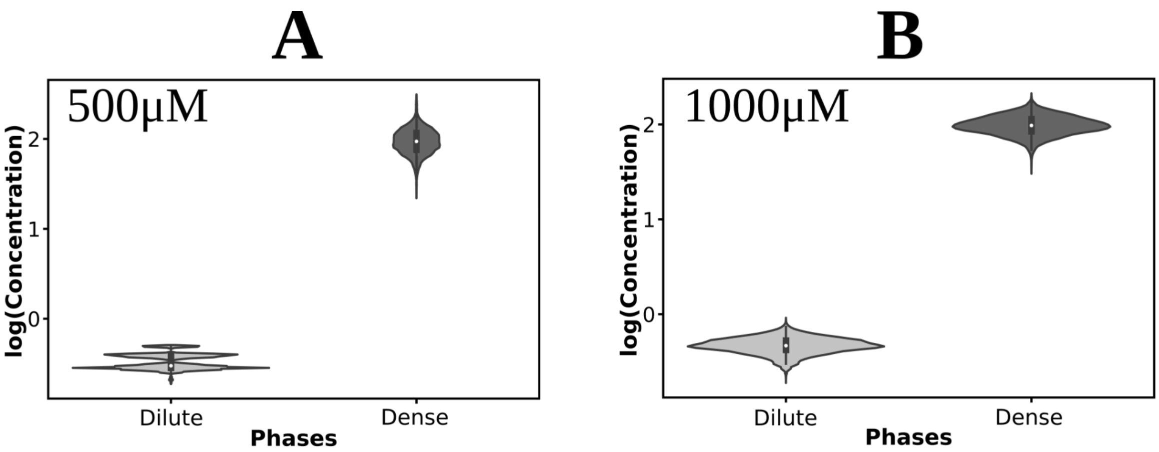


**Figure S4: LLPS-like characteristics in protein aggregates are robust across the protein concentration.** Figure A and B represent the probability distribution of protein concentrations (in the form of logarithmic function) in dilute and dense phases in violin plots for the systems containing 500 and 1000 µM of protein respectively dissolved in 50 mM aqueous NaCl solution at 300 K temperature.

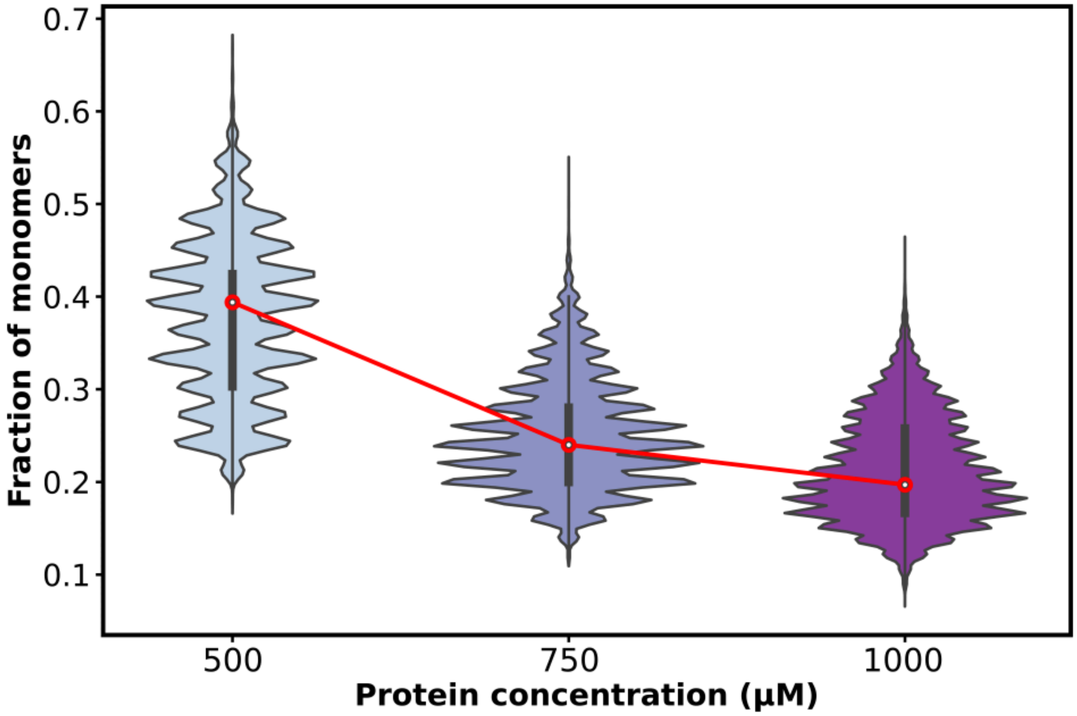


**Figure S5: Effect of variation of protein concentration on phase behavior of Aβ40 protein.** Figure shows the probability distribution (in violin plot) of fraction of number of monomers (number of monomers at an instant/total number of protein chains in the system) of Aβ40 protein in 50 mM aqueous NaCl medium at 300 K temperature for variation of protein concentration from 500 to 750 to 1000 μM.

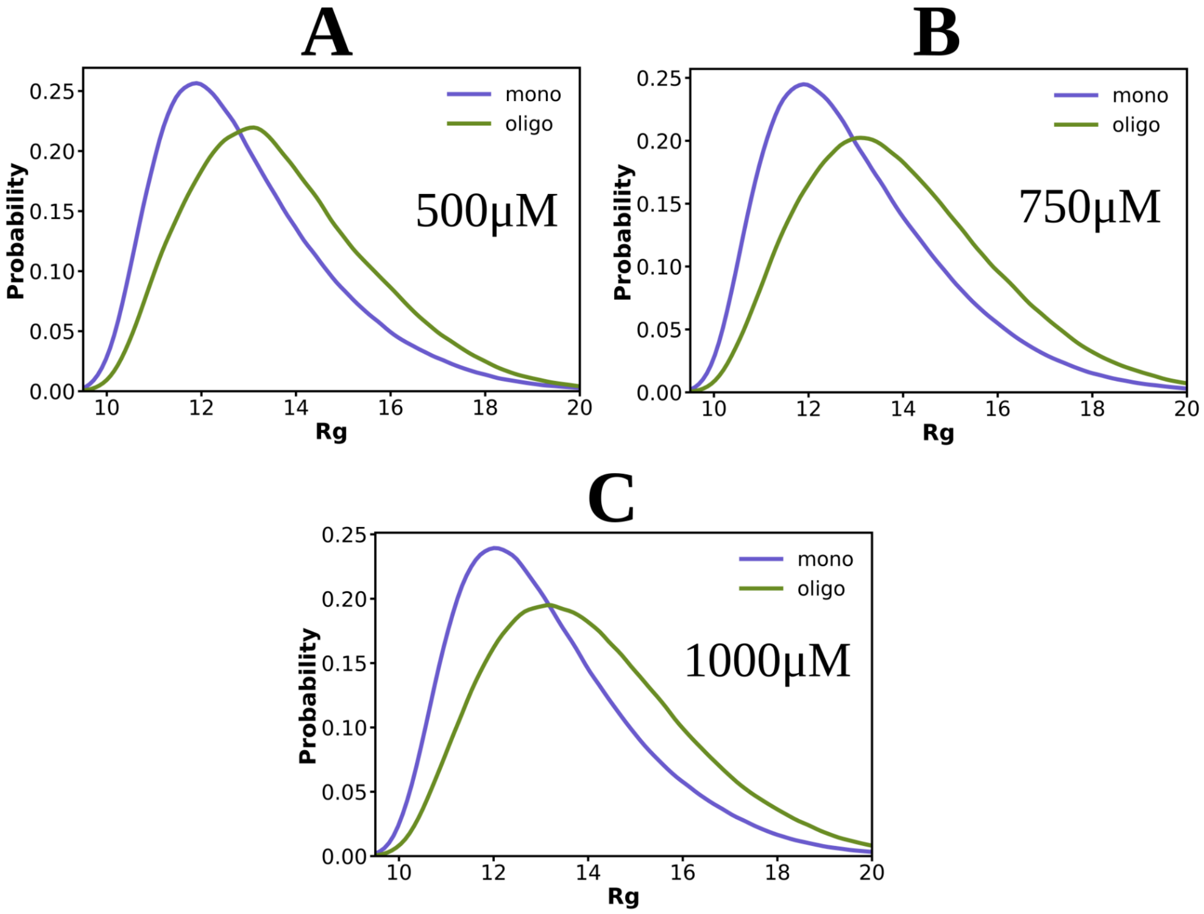


**Figure S6: Protein chain extension in oligomers remains robust across protein concentration.** Figure A, B and C represent $R_g$ of Aβ40 protein chains in monomeric and oligomeric (hexamer and beyond) state in 50 mM aqueous NaCl medium at 300 K temperature for variation of protein concentration from 500 to 750 to 1000 μM respectively.

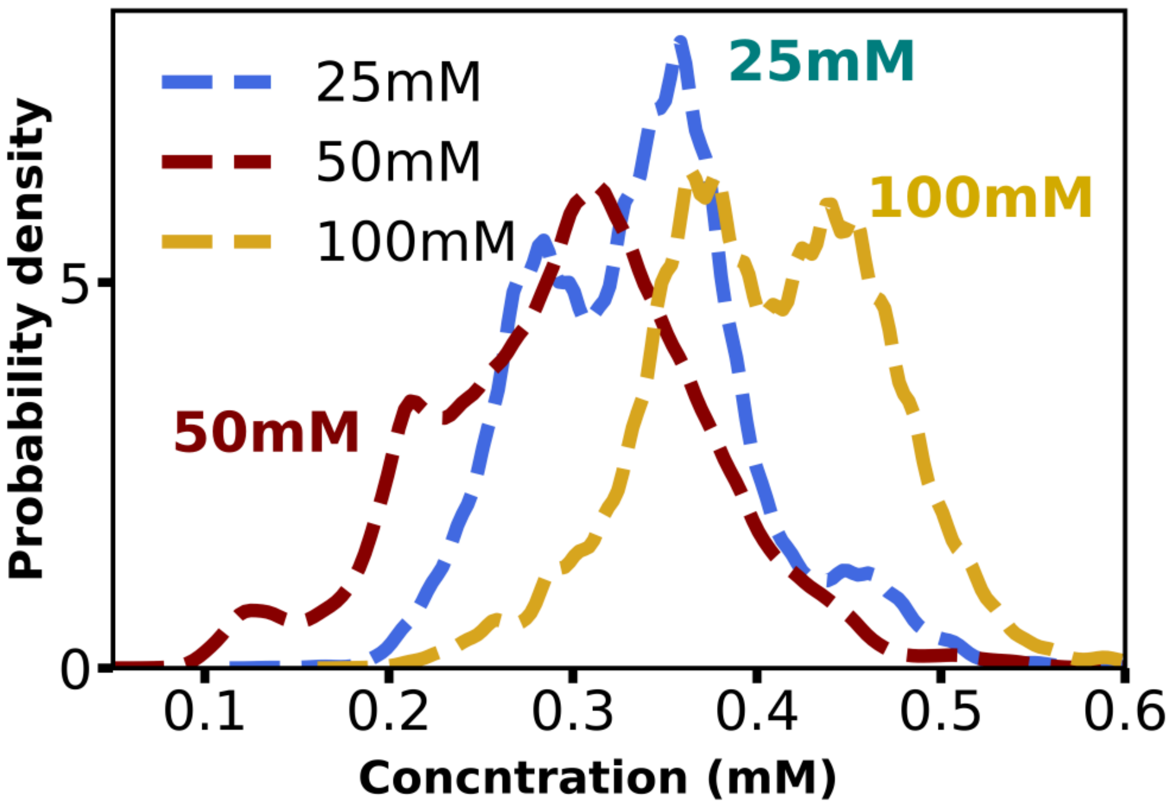


**Figure S7: Effect of changing salt concentration on protein concentration in the dilute phase.** Figure represents the change in protein concentration in the dilute phase during the process of aggregation for the system containing 750 μM Aβ40 protein at 300 K temperature on variation of salt (NaCl) concentration from 25 to 50 to 100 mM.

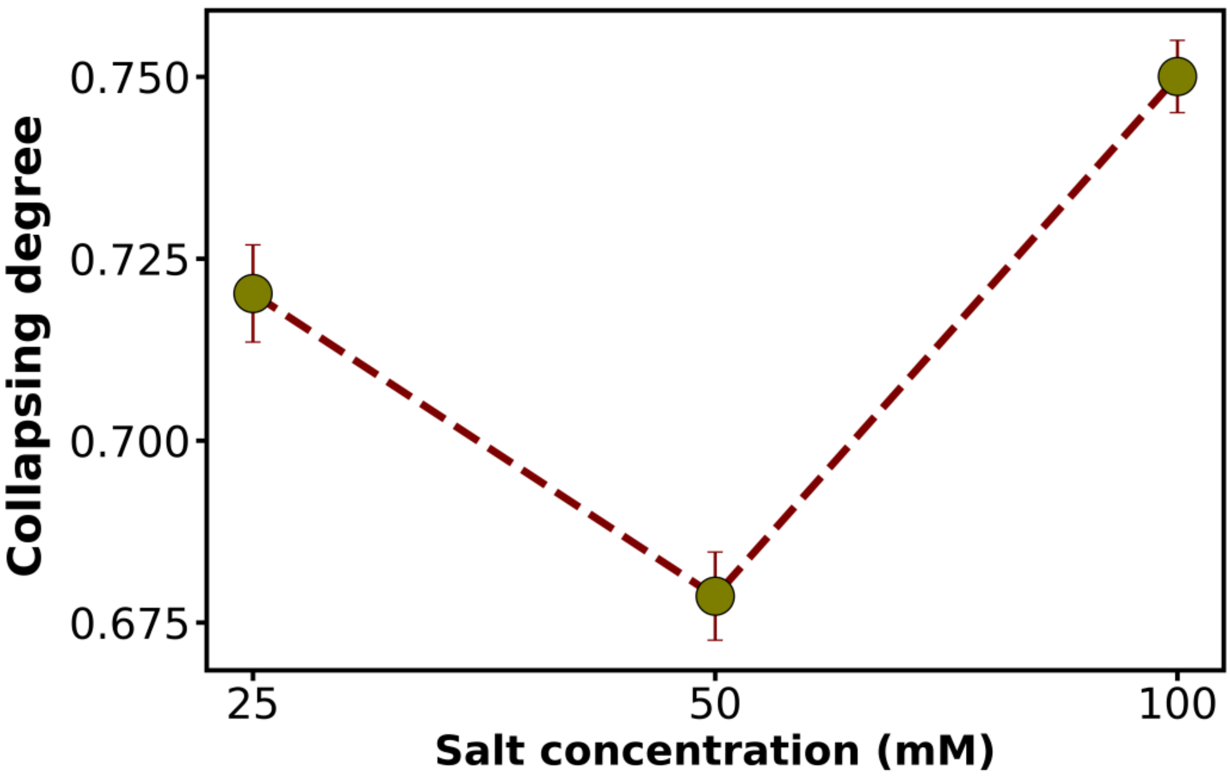


**Figure S8: Degree of collapse on changing salt concentration.** Figure shows the change of the value of collapsing degree (see maintext, equation 5) for the system containing 750 μM Aβ40 protein at 300 K temperature on variation of NaCl salt concentration from 25 to 50 to 100 mM. The error values are obtained through averaging over different replicas.

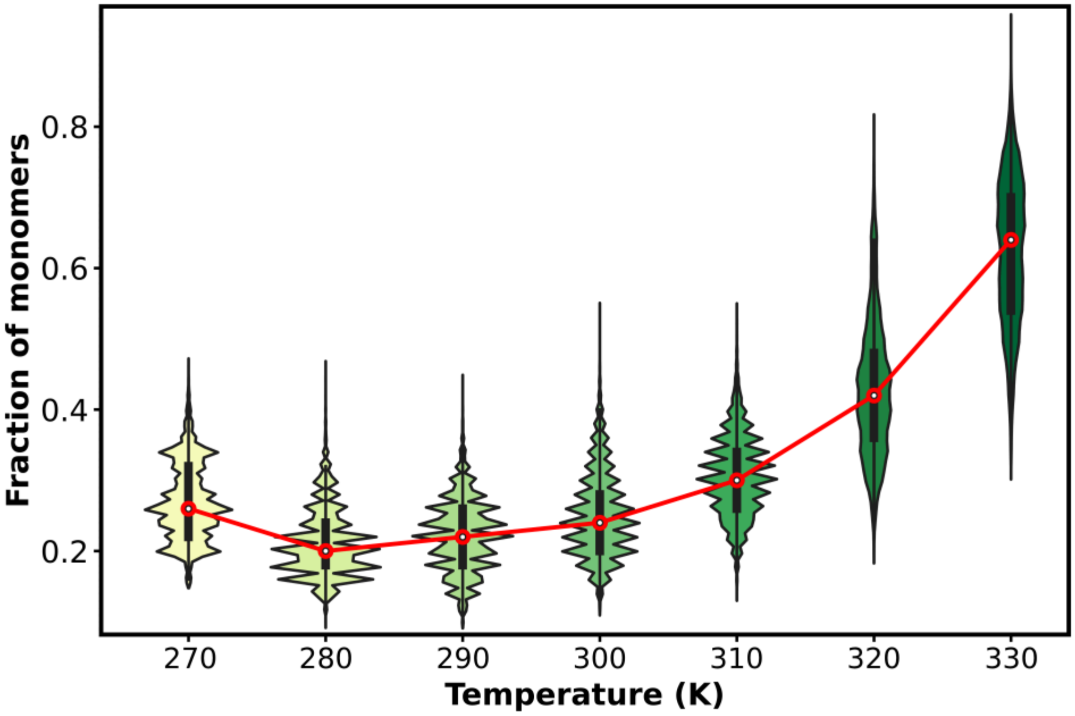


**Figure S9: The variation of fraction of protein monomers on influence of temperature.** Figure shows the fraction of monomers (the ratio of the number of protein chains in the monomeric state to the total number of protein chains in the system) of the Aβ40 protein in the systems containing 750 μM of protein concentration in 50 mM salt solution over the temperature range from 270 to 330 K with the regular interval of 10 K i.e. at 270, 280, 290, 300, 310, 320 and 330 K.

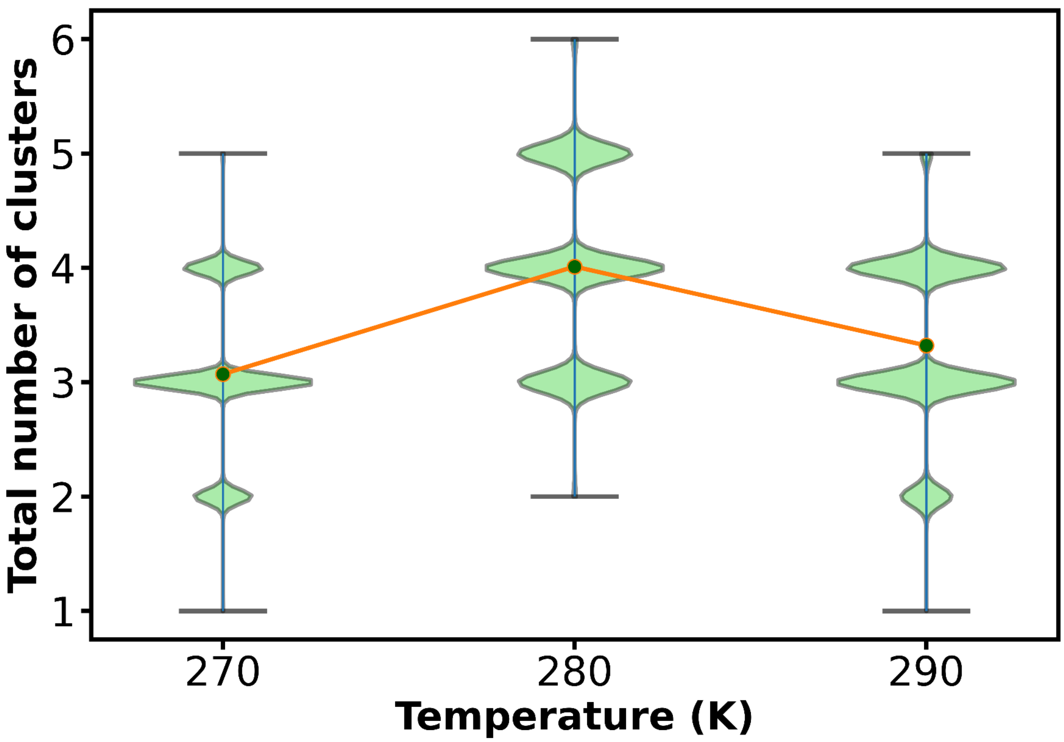


**Figure S10: Alteration of total number of clusters with temperature.** Figure shows the change of the total number of clusters formed (No. of protein chains involved in the cluster ≥ 6) for the system containing 750 μM Aβ40 protein in aqueous 50 mM NaCl solution at 270, 280 and 300 K temperature. The calculation is performed over the extended simulation trajectories corresponding to each of the systems.

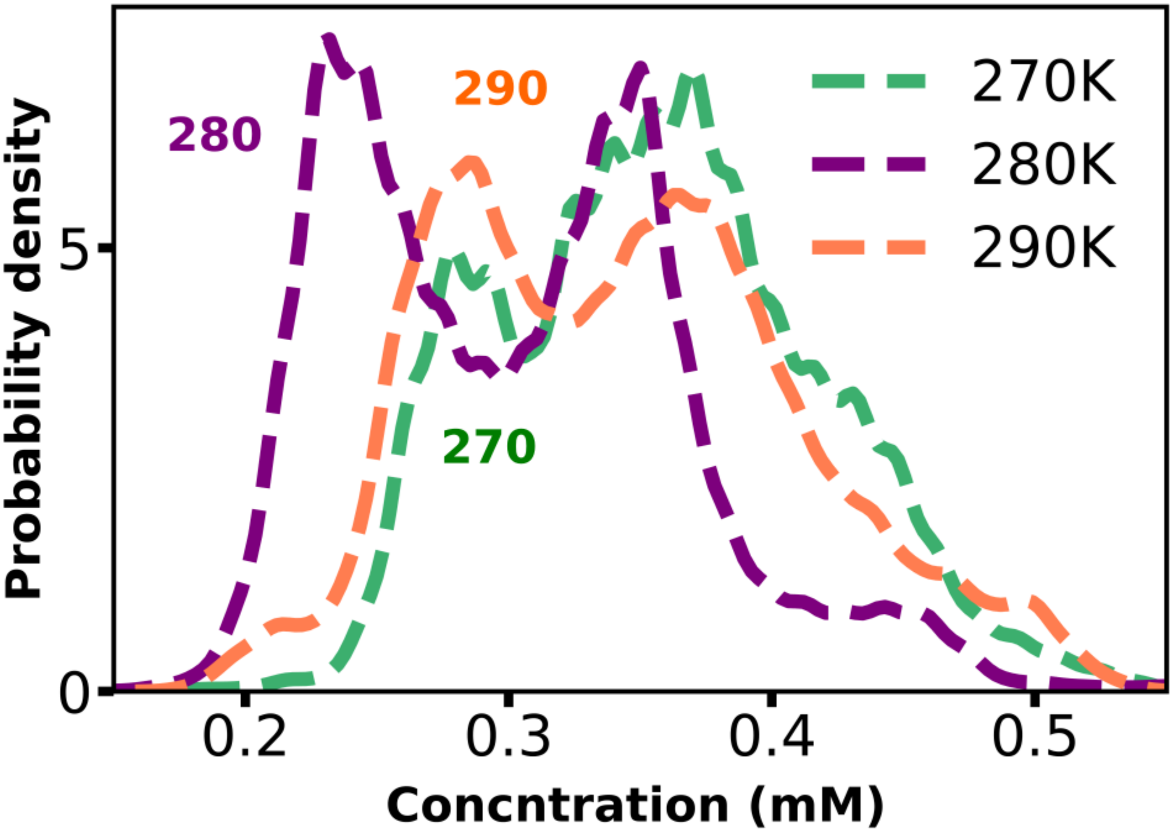


**Figure S11: Effect of changing temperature on protein concentration of the dilute phase.** Figure represents the change in protein concentration in the dilute phase during the process of aggregation for the system containing 750 µM Aβ40 protein in 50 mM NaCl salt solution for variation of temperature from 270 to 280 to 290 K. The calculation is performed over the extended simulation trajectories corresponding to each of the systems.

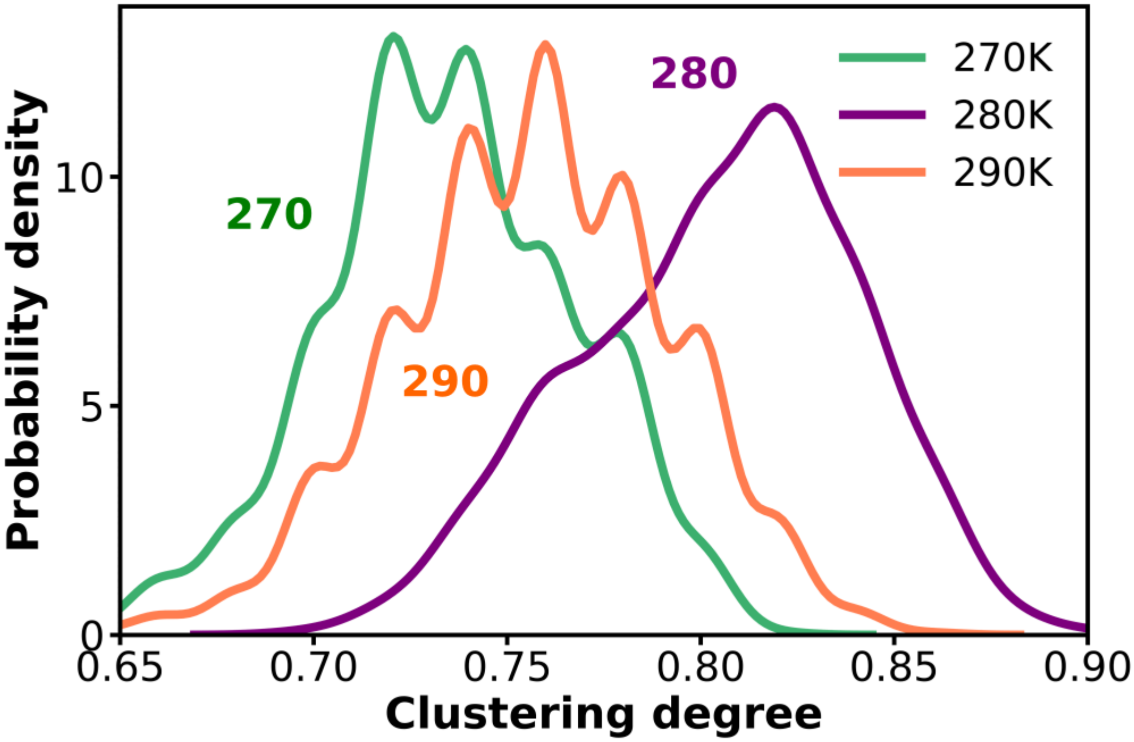


**Figure S12: Variation of clustering degree on changing solution temperature.** Figure shows the change of the value of clustering degree (see main text equation 4) for the system containing 750 μM Aβ40 protein in 50 mM NaCl salt aqueous solution on alteration of temperature from 270 to 280 to 290 K. The calculation is performed over the extended simulation trajectories corresponding to each of the systems.

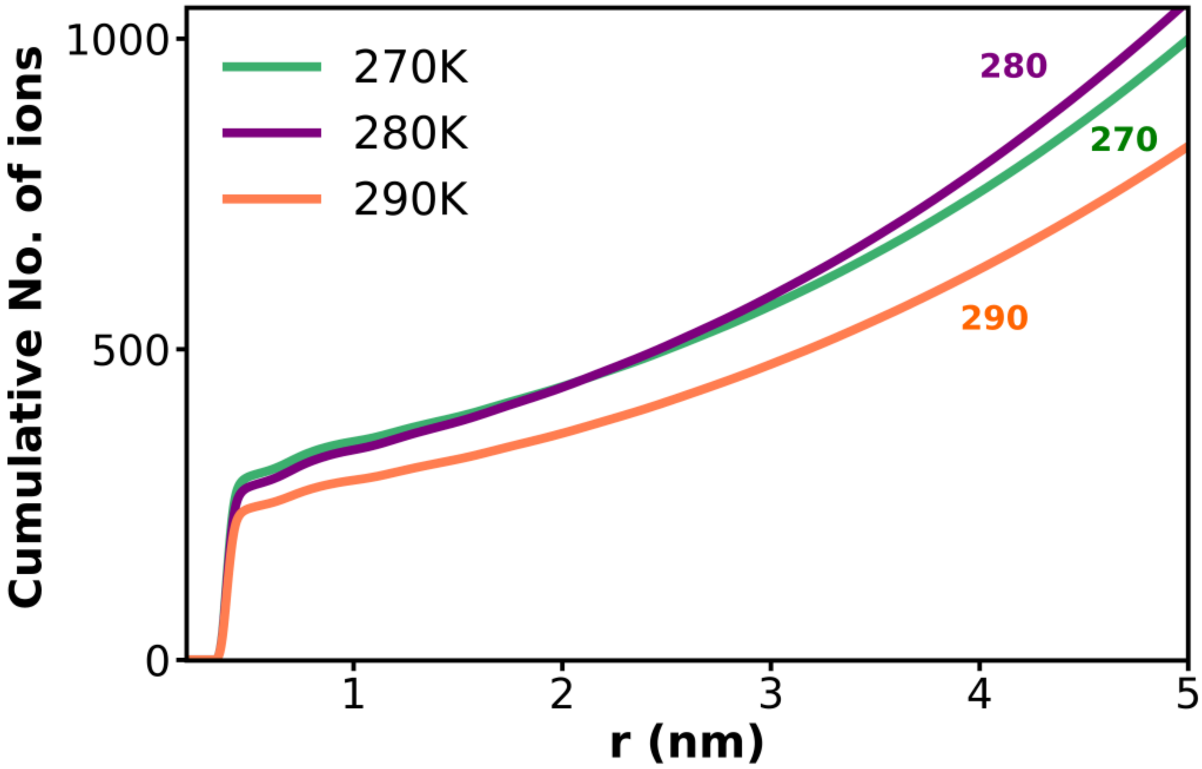


**Figure S13: Variation of number of salt ions near protein on changing solution temperature.** Figure shows the change of the number of salt species near the protein molecules for the system containing 750 μM Aβ40 protein in 50 mM NaCl salt aqueous solution on alteration of temperature from 270 to 280 to 290 K. The calculation is performed over the extended simulation trajectories corresponding to each of the systems.

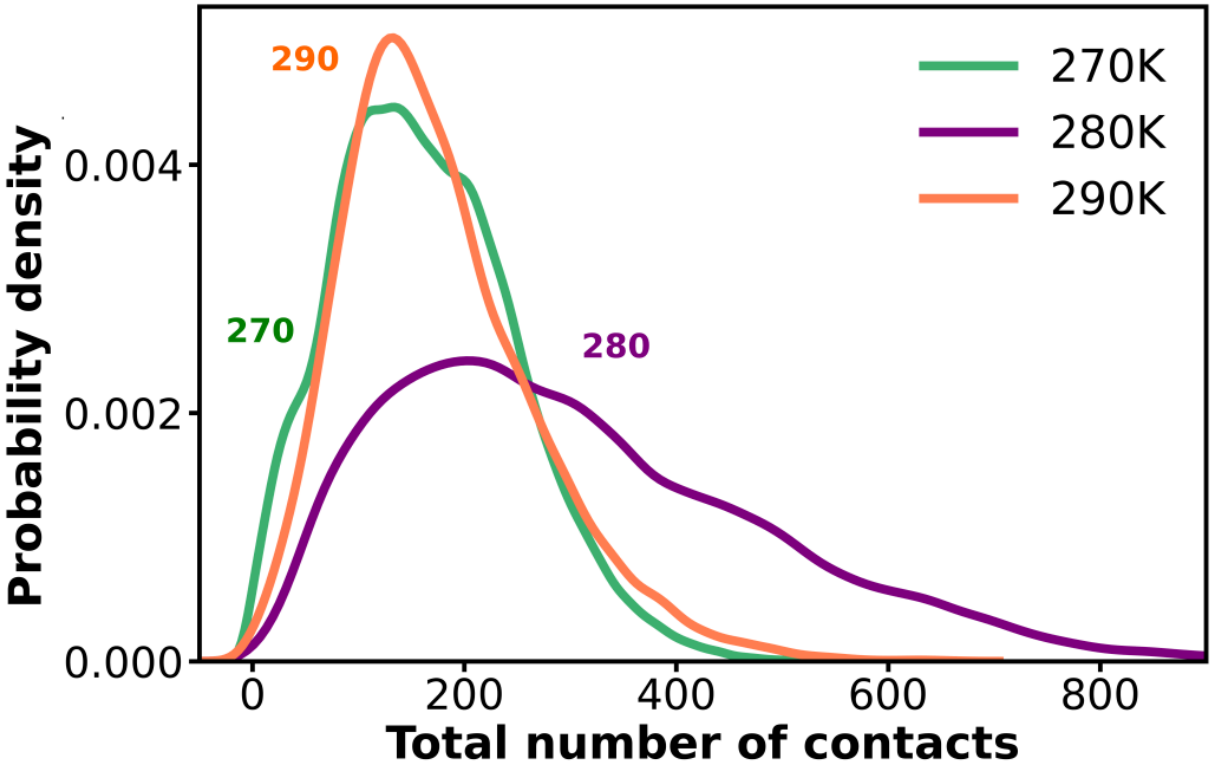


**Figure S14: Variation of total number of interprotein contacts on changing solution temperature.** Figure shows the change of the number of the total contact among the protein molecules involved in oligomers containing number of protein chains ≥ 6 for the system containing 750 μM Aβ40 protein in 50 mM NaCl salt aqueous solution on alteration of temperature from 270 to 280 to 290 K. The calculation is performed over the extended simulation trajectories corresponding to each of the systems.

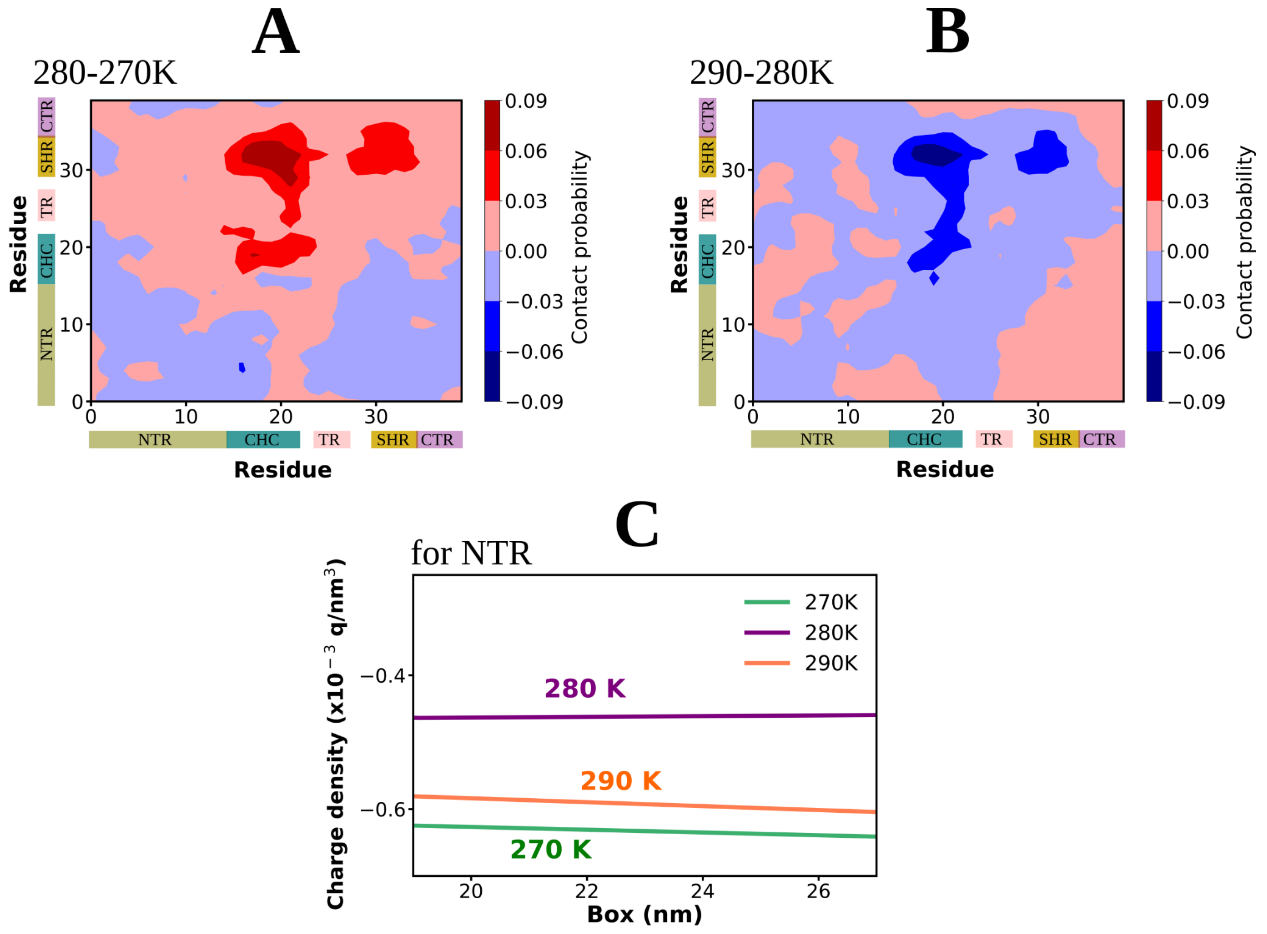


**Figure S15: Molecular insights for temperature dependent reentrant behavior of Aβ40.** Figure A and B shows the differential interchain contact maps of Aβ40 (750μM) protein chains in dense phase corresponding to the change in temperature from A. 270 to 280 K (interchain contact probability at 270 K is subtracted from that of 280 K) and B. 280 to 290 K (interchain contact probability at 280 K is subtracted from that of 290 K) in presence of 50 mM NaCl salt. The different regions of protein is represented as the N-terminal region (NTR, residues 1-16) in khaki, the central hydrophobic core (CHC, residues 17-21) in teal, the turn (TR, residues 24-27) in pink, the secondary hydrophobic region (SHR, residues 30-35) in golden, and the C-terminal region (CTR, residues 36-40) in purple color. Figure C shows the profile of charge density (see method) corresponding to the N-terminal region (NTR) of the Aβ40 protein in 50 mM aqueous NaCl solution for the temperatures of 270, 280 and 290 K. The calculation is performed over the extended simulation trajectories corresponding to each of the systems.
